# Genomic Surveillance of Respiratory Syncytial Virus among Patients with Acute Respiratory Infection through Hospital-Based Influenza Surveillance Platforms in Bangladesh, August 2024–December 2025

**DOI:** 10.64898/2026.07.02.736091

**Authors:** Md. Shaheen Alam, Yeasir Karim, Riaz Rahman Shanto, Rubel Howlader, Tanzim Rahman, Muhammad Talha, Fahmida Chowdhury, Mohammad Jubair, Mst. Noorjahan Begum, Mustafizur Rahman

## Abstract

**Background:** RSV is one of the major contributing factors of acute lower respiratory tract infection among younger children who are below five years old; over 95% of the burden falls on low- and middle-income countries. Bangladesh faces a substantial pediatric RSV burden yet lacks post-pandemic whole-genome data. Approval of nirsevimab and maternal vaccines (Arexvy, Abrysvo), underscores the need for region-specific genomic surveillance to inform prevention and control strategies.

**Objectives:** To conduct the whole-genome sequencing (WGS) study of RSV in Bangladeshi population, characterizing viral genotypic diversity, phylogenetics, F and G protein mutations, glycosylation dynamics, while generating and depositing complete genomes in GISAID.

**Methods:** Between August 2024 and December 2025, of 1,390 RSV-positive specimens (Ct ≤35), 59 high-viral-load samples (Ct ≤25) were selected for whole-genome sequencing using Oxford Nanopore Technology (ARTIC primers), assembled with MIRA v2.0.0, and analyzed via Augur/Nextstrain, Nextclade, and NetNGlyc/NetOGlyc 4.0.

**Results:** Of 11,874 patients, 1,390 (11.7%) were RSV-positive (RSV-A 94.6%). From selected 59 RSV-positive cases, 49 high-quality genomes were generated (83.1% pass; 94.7% completeness; median depth 1,500×, range 443-4,948): 43 RSV-A (ON1; A.D.3.7 63%, A.D.3 19%, A.D.3.12 9%, A.D.3.1 7%, A.D.1.11 2%) and 6 RSV-B (BA9/B.D.E.1). S276N at antigenic site II (34.9% RSV-A) and S389P in all RSV-B were detected; neither confers resistance to nirsevimab or palivizumab. An F protein N75 N-glycosylation site was fixed in all RSV-A; RSV-B acquired HVR2 N256 glycan in 67%. All 49 genomes were deposited in GISAID.

**Conclusion:** This WGS study of RSV in the Bangladeshi population confirms LMIC nanopore surveillance feasibility, multi-lineage co-circulation, and intact conservation of all vaccine and antibody targets. Progressive glycan remodeling warrants monitoring. These findings establish a genomic baseline to guide nirsevimab and maternal vaccine deployment in Bangladesh, with direct relevance for RSV surveillance programs across South Asia.

## 1. Background

Respiratory syncytial virus (RSV) is one of the major contributing factors for acute lower respiratory tract infection (ALRI) among younger children who are below five years globally, responsible for approximately 33 million cases and more than 100,000 deaths per year, including above 95% of this burden within low- and middle-income countries (LMICs)[1]. South Asia bears the highest absolute RSV mortality, and the 2022–2023 approval of nirsevimab and maternal RSV vaccines (Arexvy, Abrysvo) has made region-specific genomic surveillance a high-priority need[2]. RSV circulates as two major subgroups- RSV-A (predominantly ON1 genotype) and RSV-B (predominantly BA genotype)- with the F protein being the sole target of all licensed interventions and the G protein the principal driver of genotypic diversification[3, 4].

In Bangladesh, RSV is a major pediatric emergency. National SARI sentinel surveillance recorded an average RSV positivity of 49% during October 2022–March 2023 [6], and a 2025 Lancet Global Health study from the Child Health Research Foundation (CHRF) documented marked seasonal peaks with substantial healthcare-resource implications, estimating that RSV alone occupied up to 20% of bed capacity in Bangladesh’s largest pediatric hospital during the 2019 season [5]. A prospective community cohort (August 2021–June 2023) in rural Bangladesh measured RSV-associated ARI cases at a rate of 107.7 per 1,000 children per year, peaking at 37.6% positivity in August–September [6], while RSV accounted for 12–13% of under-five SARI deaths, predominantly in infants below six months, both before and during the COVID-19 pandemic [7]. Molecularly, the only prior characterization was a partial G-gene study (2008–2012) identifying RSV-A (82%) and ON1/BA lineages as dominant [8], leaving a critical post-pandemic whole-genome sequencing (WGS) gap for Bangladesh.

Across countries bordering Bangladesh, RSV genomic data remain sparse. In India, partial G-gene studies documented co-circulating ON1/GA2 (RSV-A) and BA/GB5 (RSV-B) genotypes, with alternating subgroup dominance across seasons [9, 10]. Pakistan has reported multiple RSV genotypes in children — NA1/GA2 for RSV-A, along with BA9/BA10 in case of RSV-B, and a recent Islamabad outbreak WGS study (2022–2023) revealed ON1 and BA-10 lineages with unique F protein substitutions [11, 12]. A comprehensive phylogenetic and phylodynamic analysis from Karachi [13] further characterized RSV-A along with RSV-B evolutionary dynamics in children under five, while infant mortality surveillance in Karachi documented high RSV positivity with monsoon-season peaks[13, 14]. In Myanmar (eastern border), RSV-A A.D.3 persisted through the pandemic with a major RSV-B (B.D.E.1) resurgence in 2023[15]. Nepal reported a high community RSV-ARI incidence of 213 per 1,000 infant-person-years, RSV-B predominance in 69.4% of hospitalized LRTI cases in 2023, and significant cost-of-illness burden [16, 17]. In Bhutan, sentinel surveillance found children under two years at three-fold higher odds of RSV positivity among SARI admissions[18]. Together, these data highlight RSV as a shared regional threat, yet the absence of WGS data across this contiguous geographic zone severely limits cross-border phylodynamic inference.

To address this gap, we leveraged the icddr,b Hospital-Based Influenza Surveillance (HBIS) platform to perform the first WGS study of RSV in the Bangladeshi population. Our objectives were to: (i) determine genotypic and sub-lineage diversity of co-circulating RSV-A and RSV-B; (ii) establish international phylogenetic relatedness of local lineages; (iii) characterize the mutational landscape of the fusion(F) and attachment glycoprotein (G) proteins; (iv) profile N-and O-glycosylation dynamics as immune evasion markers; and (v) deposit complete genomes into GISAID to strengthen regional genomic surveillance.

## 2. Method

### 2.1. Study Design, Setting, and Population

This investigation was conducted within the framework of Bangladesh’s existing hospital-based influenza surveillance platform, which has been operational for several years. The surveillance system systematically collects clinical specimens from patients presenting to sentinel hospitals with acute respiratory symptoms meeting the World Health Organization’s case definitions for severe acute respiratory infection (SARI) or influenza-like illness (ILI).

Between August 2024 and December 2025, a total of 11,874 patients presenting with acute respiratory symptoms were enrolled across participating sentinel hospitals. Of these, 7,955 (67.0%) met the SARI case definition, while 3,965 (33.4%) met the ILI case definition. The study population encompassed a broad age range, with particular emphasis on children under five years of age, given their documented vulnerability to severe respiratory viral infections.

For each enrolled participant, field staff collected paired nasopharyngeal and oropharyngeal swabs, which were placed into viral transport medium and immediately maintained under cold chain conditions at -70°C or lower. Specimens were subsequently transported to the central laboratory at icddr,b for long-term cryopreservation and molecular analysis. Categorical variables were compared using Pearson’s chi-square test (χ²); a p-value < 0.05 was considered statistically significant. Continuous variables are reported as mean ± standard deviation (SD) or median with interquartile range (IQR) and range, as appropriate. Mutation frequencies are expressed as counts and proportions of the total sequences per subtype. All statistical analyses and data visualizations were performed in R (version 4.5.1; R Foundation for Statistical Computing, Vienna, Austria) using the ggplot2 and dplyr packages. The study design and our analysis are shown in Figure 1.

**Figure 1.**
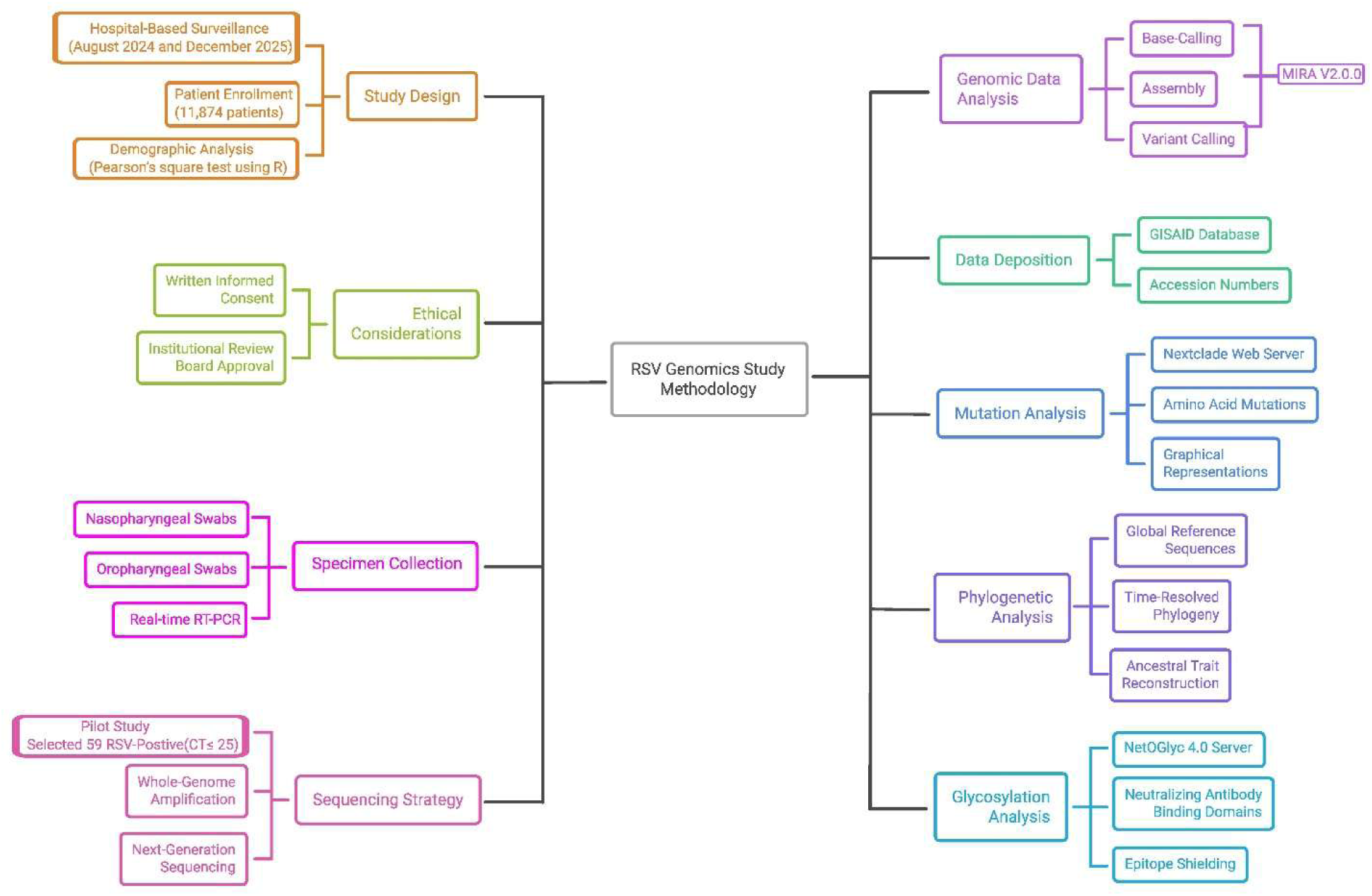
Schematic workflow of the RSV whole-genome sequencing study. The flowchart illustrates the stepwise study design, from patient enrollment at sentinel hospitals (August 2024–December 2025; n = 11,874) through rRT-PCR screening, subtyping, high-viral-load specimen selection (Ct ≤ 25), Oxford Nanopore Technology sequencing, MIRA pipeline quality control, and downstream phylogenetic, mutational, and glycosylation analyses. Numbers at each stage indicate the sample counts retained or excluded at that step.

### 2.2. Specimen Collection and RSV Detection using Real-time RT-PCR

Detailed procedures for specimen collection, RNA extraction, and real-time RT-PCR screening (including the Ct threshold used for positivity) have been previously described (2). For samples positive for RSV, subtyping into RSV-A and RSV-B was performed using the protocol established by Fernanda de Paris et al. (3). A sample was considered RSV-positive if the cycle threshold (Ct) value was ≤ 35. CT values were recorded for all positive specimens.

### 2.3. Strategy for Selecting Sequencing Samples

Before large-scale whole-genome sequencing, a pilot feasibility study was conducted on a representative subset of 59 RSV-positive clinical specimens. To ensure optimal sequencing performance characterized by robust consensus genome assembly, high-depth coverage, and minimal host nucleic acid contamination, selection was restricted to high-titer specimens defined by rRT-PCR Ct values ≤ 25.

### 2.4. HRSV Whole-Genome Amplification

First-strand cDNA was synthesized from RSV-positive samples with a Ct value ≤25 following the PrimeScript™ 1st Strand cDNA Synthesis Kit (Takara Bio Inc., Cat. #6110A). Briefly, a 13 µL mixture containing 1 µL dNTP mix, 1 µL random primers, 7 µL template RNA, and 4 µL RNase-free water was incubated at 65 °C for 5 minutes for RNA denaturation and then placed on ice. Subsequently, 7 µL of reverse transcription master mix (4 µL 5× buffer, 1 µL 0.1 M DTT, 1 µL reverse transcriptase, 1 µL RNase-free water) was added. The final 20 µL reaction was incubated at 25 °C for 2 minutes, 50 °C for 60 minutes, and 70 °C for 15 minutes. cDNA was stored at - 20°C. Conventional PCR for whole-genome amplification was performed using GoTaq® Hot Start Polymerase (Promega). For RSV-A, primers and cycling conditions followed Wang L. et al. (4). For RSV-B, the ARTIC Network-designed primer set (version 1) was used (5). Primer pools were prepared separately for RSV-A (20 primers) and RSV-B (101 primers), with odd- and even-numbered primers in separate pools. Each 25 µL PCR contained 5X GoTaq Flexi Buffer, 2.5 mM MgCl₂, 0.2 mM each dNTP, 0.015 µM final concentration of each primer, and 3.5 µL cDNA. Separate master mixes were prepared for odd and even primer pools. Cycling conditions for RSV-A were: 95°C for 2 min; 35 cycles of 98°C for 15 sec and 63°C for 7 min; hold at 4°C. For RSV-B: 95°C for 2 min; 35 cycles of 98°C for 15 sec and 63°C for 5 min; final extension at 68°C for 5 min; hold at 4°C.

### 2.5. Next-Generation Sequencing (NGS) using Oxford Nanopore Technology

PCR products from odd and even pools were visualized on a 1.5% agarose gel. For each sample, products from both pools were combined (40 µL total) and purified using a 0.8× bead cleanup (HighPrep™ PCR Clean-up System, MagBio). The eluted DNA (in 20 µL nuclease-free water) was quantified using the Qubit dsDNA BR Assay Kit (Thermo Fisher). Sequencing libraries were prepared using the Ligation Sequencing Amplicons – Native Barcoding Kit 96 V14 (SQK-NBD114.96, Oxford Nanopore Technologies) per the manufacturer’s protocol. The final pooled library was loaded onto a FLO-MIN114 (R10.4.1) flow cell and sequenced on a Mk1C device for approximately 30 hours.

### 2.6. Genomic Data Analysis and Bioinformatics

Raw FAST5 data were base-called and demultiplexed using Guppy (v6.5.7) on the Mk1C device (minimum Q-score: 8). Resulting FASTQ files were quality-checked and assembled using MIRA v2 (part of the CDC’s Viral NGS pipeline) for reference-based mapping and consensus generation. References used were RSV-A (KY654518) and RSV-B (KF640637). The MIRA pipeline (v2.0.0, module ont-rsv) performed assembly, variant calling, and automated quality control. A sample “passed” if it achieved a median coverage ≥50x, covered ≥90% of the reference genome, and contained fewer than 20 minor single-nucleotide variants (SNVs) with frequency ≥5%. Samples failing these criteria were excluded.

### 2.7. Phylogenetic and Evolutionary Analysis

To contextualize the Bangladeshi sequences, we obtained global reference sequences from GISAID. For RSV-A, 17,773 sequences were retrieved, from which 5% (n=888) were randomly selected and combined with our study sequences. For RSV-B, 19,557 sequences were retrieved, with 5% (n=977) randomly selected and combined with our study sequences. Phylogenetic analysis was conducted using the Augur toolkit within a Snakemake workflow. Sequences were aligned to reference genomes (RSV-A: hRSV/A/England/397/2017, PP109421; RSV-B: hRSV/B/Australia/VIC-RCH056/2019, OP975389) as suggested by Nextclade. A time-resolved phylogeny was inferred using a coalescent model with marginal date estimation, filtering molecular clock outliers (IQD=4). Ancestral trait reconstruction (geography, mutations) was performed via joint inference. The final phylogeny was visualized in Auspice.

### 2.8. Mutation Analysis

FASTA files of consensus genomes were analyzed using the Nextclade web server. Amino acid mutations across the entire genome (relative to the aforementioned references) were identified and extracted. Subsequent graphical representations of mutations were generated using R.

### 2.9. N-glycosylation and O-glycosylation

RSV F and G protein sequences were analyzed for potential N-linked glycosylation using the NetNGlyc 1.0 server (threshold ≥ 0.5). N-X-S/T sequons meeting the threshold were considered high-confidence glycosylation sites. Residues identified as biologically inaccessible (e.g., cytoplasmic tail positions) were noted but excluded from immunological interpretation. RSV F protein sequences were additionally analyzed for potential O-linked glycosylation using the NetOGlyc 4.0 server (G-score threshold > 0.5). Only residues within the mature protein coding region (residues 6–574) were considered high-confidence hits. Predicted sites were mapped to known neutralizing antibody binding domains defined by the following coordinates: Site Ø (61–69, 196–212), Site V (160–180), Site II (255–275), Site I (382–396), and Site IV (422–438). Data processing, statistical quantification of epitope shielding, and visualization were performed using **R** with the ggplot2 and dplyr packages.

### 2.10. Data Deposition

All 49 complete RSV genome sequences generated in this study have been deposited in GISAID (https://www.gisaid.org/) under accession numbers EPI_ISL_20094933, EPI_ISL_20095115–20095119, EPI_ISL_20212052–20212064, and EPI_ISL_20287914–20287943.

### 2.11. Ethical Considerations

Respiratory specimens and associated clinical and demographic data were collected through the ongoing HBIS platform under protocol **PR-22150**, with written informed consent obtained from adult participants or parents/legal guardians of participating children. The present RSV genomics study was conducted under the **PR**-24122, which was validated by the Institutional Review Board (IRB) of icddr,b, and utilized archived HBIS specimens and metadata for genomic analysis.

## 3. Results

### 3.1. Demographic Characteristics and Temporal Trends

Over the 17-month surveillance period, 11,874 illness episodes were documented from 11,872 unique individuals. Of these, 7,955 (67.0%) were classified as SARI and 3,965 (33.4%) as ILI, with RSV positivity confirmed in 1,390 episodes (11.7%). As shown in Table 1, male predominance was observed in both groups (SARI: 59%, ILI: 63%; p < 0.001), and age distribution differed significantly (p < 0.001). Infants aged 6–11 months represented the largest SARI subgroup (1,779 cases, 22%), while adults aged 18–39 years comprised the majority of ILI cases (1,498 cases, 38%). Hospital-level variation was also significant (p < 0.001), with Chittagong Medical College Hospital managing the highest proportion of SARI cases (16%) and Jashore 250-bed General Hospital treating the most ILI cases (20%), likely reflecting local referral patterns. Demographic characteristics are summarized in **Table 1**.

**Table 1.** Demographic distribution of SARI and ILI episodes.

| Characteristics | SARI<sup>1</sup> | ILI<sup>1</sup> | Overall<sup>1</sup> | p-value<sup>2</sup> |
| --- | --- | --- | --- | --- |
| **Sex** |  |  |  | <0.001 |
| Female | 1,603 (41 %) | 2,962 (37 %) | 4,565 (38 %) |  |
| Male | 2,316 (59 %) | 4,993 (63 %) | 7,309 (62 %) |  |
| **Age group** |  |  |  | <0.001 |
| <2 months | 45 (1.1 %) | 370 (4.7 %) | 415 (3.5 %) |  |
| 2-5 months | 184 (4.7 %) | 1,233 (16 %) | 1,417 (12 %) |  |
| 6-11 months | 369 (9.4 %) | 1,779 (22 %) | 2,148 (18 %) |  |
| 12-23 months | 36 (0.9 %) | 190 (2.4 %) | 226 (1.9 %) |  |
| 24-59 months | 154 (3.9 %) | 331 (4.2 %) | 485 (4.1 %) |  |
| 5-17 years | 946 (24 %) | 1,172 (15 %) | 2,118 (18 %) |  |
| 18-39 years | 1,498 (38 %) | 1,295 (16 %) | 2,793 (24 %) |  |
| 40-59 years | 525 (13 %) | 819 (10 %) | 1,344 (11 %) |  |
| ≥60 years | 162 (4.1 %) | 765 (9.6 %) | 927 (7.8 %) |  |
| **Hospital** |  |  |  | <0.001 |
| Chittagong Medical College Hospital | 816 (21 %) | 1,277 (16 %) | 2,093 (18 %) |  |
| Cumilla Medical College Hospital | 126 (3.2 %) | 970 (12 %) | 1,096 (9.2 %) |  |
| Dinajpur Medical College Hospital | 749 (19 %) | 760 (9.6 %) | 1,509 (13 %) |  |
| Jahurul Islam Medical College | 651 (17 %) | 947 (12 %) | 1,598 (13 %) |  |
| Jalalabad Ragib-Rabeya Medical College Hospital | 727 (19 %) | 942 (12 %) | 1,669 (14 %) |  |
| Jashore 250-bed General Hospital | 785 (20 %) | 769 (9.7 %) | 1,554 (13 %) |  |
| Khulna Medical College Hospital | 62 (1.6 %) | 605 (7.6 %) | 667 (5.6 %) |  |
| Rajshahi Medical College Hospital | 1 (<0.1 %) | 743 (9.3 %) | 744 (6.3 %) |  |
| Sher-e-Bangla Medical College Hospital | 2 (<0.1 %) | 942 (12 %) | 944 (8.0 %) |  |

<sup>1</sup>Values represented as n (%).
<sup>2</sup>Pearson's Chi-square test ( $\chi^2$ ). $\chi^2 = \sum [(Observed - Expected)^2 / Expected]$ . A p-value < 0.05 was considered statistically significant.

Monthly trends (Supplementary Figure 1) revealed distinct seasonal periodicity for SARI, with peaks during August–October 2024 and July–September 2025, while ILI cases followed a similar but lower pattern (190–270 cases monthly). RSV positivity rates (Supplementary Figure 2) showed two discrete waves: a major peak in September–October 2024 (41%), a marked decline to 0–2% between May–July 2025, and a progressive resurgence from August 2025 (2%) to December 2025 (10%). This biennial pattern—characterized by a major epidemic peak followed by an interepidemic period of low circulation and subsequent resurgence—aligns with RSV epidemiology documented in Bangladesh, where studies have reported peak RSV activity during the post-monsoon months of September through December [1,2].

Age-specific analysis (Supplementary Figure 3) confirmed that children under five years bore the highest RSV burden (4,266 episodes, 36%), with infants aged 6–11 months being most vulnerable (1,779 SARI, 369 ILI) and neonates under two months contributing 370 SARI cases (5% of all SARI). These findings are consistent with previous community-based prospective studies in rural Bangladesh, which reported that the incidence of RSV-related ARI was highest during the initial six months of life at 164.5 per 1,000 child-years [3].

Monthly age-group distribution (Supplementary Figure 4) revealed that RSV transmission was consistently driven by children under five years, particularly the 6–11 months and 2–5 months groups, while adults aged ≥18 years exhibited persistently low activity (rarely exceeding 10 cases monthly). CT value analysis (Supplementary Figure 5) showed that most positive specimens fell within the ≤35 range, with a peak concentration at 20–25, and that young children demonstrated a slightly broader distribution. Finally, RSV subtyping (Supplementary Figure 6) revealed overwhelming RSV-A predominance across all age groups, with RSV-B detected only sporadically, indicating that subtype-specific susceptibility does not vary by age.

### 3.2. Laboratory Testing and RSV Characterization Results

Of the 11,874 specimens tested by rRT-PCR, 1,390 (11.7%) were confirmed positive for RSV based on a Ct threshold of ≤ 35. Among these, RSV-A accounted for 1,315 cases (94.6%), while RSV-B accounted for 71 cases (5.1%). Four cases (0.3%) were RSV-positive but could not be subtyped. The distribution of Ct values among RSV-positive cases is shown in the Supplementary Figure 5, stratified by age group (under 5 years vs. 5 years and above). Lower Ct values, indicating higher viral loads, were more frequently observed in the under-5 age group compared to older individuals. The red dashed line indicates the Ct ≤ 35 positivity threshold.

### 3.3. Outcomes of the Sample Selection Strategy

A pilot sequencing was conducted on 59 RSV-positive specimens with Ct values ≤ 25, including 50 SARI (84.7%) and 9 ILI (15.3%) cases, with a male predominance (61.0%). The median age was 0.8 years (IQR: 0.3–2.0), and 71.2% occurred in children under 1 year; only three adult cases (5.1%) were recorded. Subtype A accounted for 88.1% (52/59) and subtype B for 11.9% (7/59), with subtype A predominant in both SARI (88.0%) and ILI (88.9%) groups. Mean Ct values were 21.43 (±2.91) for RSV-A and 20.83 (±4.41) for RSV-B, with lower Ct values observed in infants aged <6 months. Despite the selection of high-titer specimens (Ct ≤ 25), 49 of the 59 samples passed quality control.

### 3.4 Genome Generation and Data Quality

A total of 59 RSV-positive nasopharyngeal swabs with high viral load (Ct ≤ 25) were selected for sequencing. Using Oxford Nanopore Technology and the ARTIC primer scheme, we generated 49 high-quality whole-genome sequences, achieving a mean reference coverage of 94.7% and a median sequencing depth of 1,500× (range: 443×–4,948×). The dataset comprised 43 RSV-A and 6 RSV-B genomes.

Using the MIRA-v2.0.0 ont-rsv module, we processed 59 samples in total; 49 (83.1%) of them passed every quality control test (Figure 2). Excessive minor variation numbers or inadequate genome coverage (<90%) were the main reasons for the failure of the ten samples. Sequencing data quality was strong among the successful samples, with an average of 32,879 total reads per sample (derived from the arithmetic mean of total reads for the 49 passed samples) and 29,957 reads mapping to the reference (derived from the arithmetic mean of mapped reads for the 49 passed samples). With a mean reference coverage of 94.7% (derived from the arithmetic mean of reference coverage percentages for the 49 passed samples) and a median coverage depth of 1,500X (derived from the arithmetic mean of the median coverage depths for the 49 passed samples) (range: 443X – 4,948X), these samples demonstrated a high degree of genome completeness (Figure 3). Of the 49 passed genomes, 6 (12.2%) were classified as RSV-B and 43 (87.8%) mapped to the RSV-A reference.

**Figure 2.**
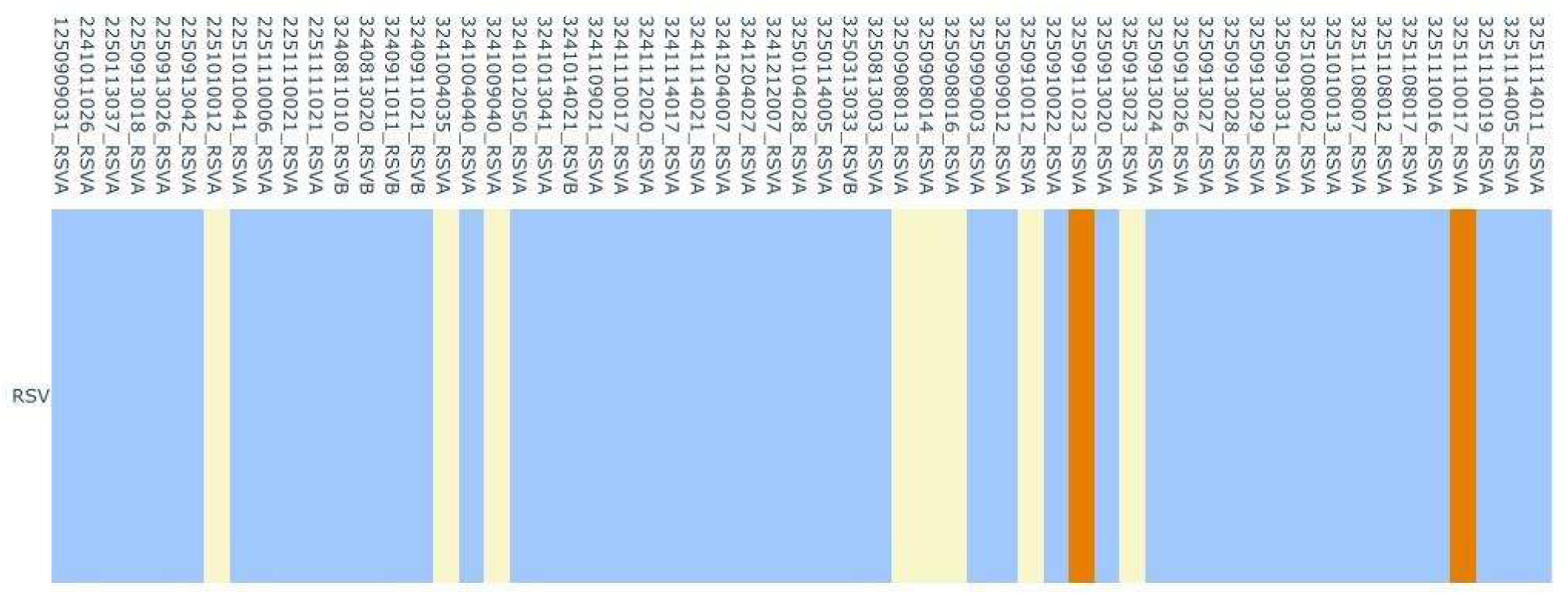
Sequencing quality control overview of the 59 processed RSV samples. The figure illustrates the proportion of samples that met all quality control (QC) criteria versus those that failed. Samples successfully passing the MIRA pipeline are indicated in **sky blue** (n=49), while samples failing due to low coverage (light yellow) or excess variants (Orange) are indicated in (n=10).

**Figure 3.**
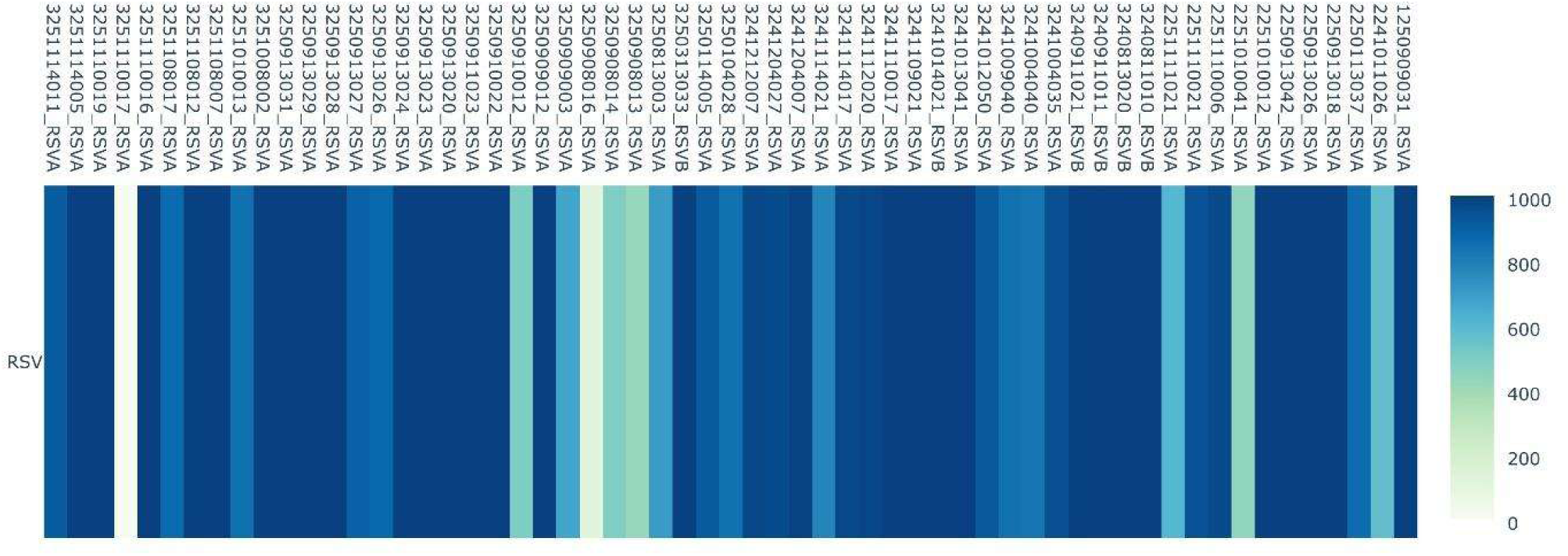
RSV genome sequencing depth across samples is displayed in a heatmap. A single RSV-positive sample (RSV-A and RSV-B) is represented by each column, and the color intensity, which ranges from low (light) to high (dark blue), depicts the mean read depth throughout the genome.

The MIRA v2.0.0 ont-rsv module rated all 49 samples as “Pass,” indicating strong sequencing performance and readiness for subsequent genomic and phylogenetic analysis, if they satisfied predetermined quality requirements. Conversely, ten samples, or 16.9% of the cohort, did not satisfy the quality control requirements. Eight samples had less than 90% reference coverage, indicating that inadequate genome coverage was the main reason for failure. The majority of these samples had reasonable read counts but inadequate coverage breadth (mean % reference covered: ∼89.7%), with one sample failing due to an exceptionally low sequencing yield (12 total reads, 1.17% coverage). Despite having sufficient coverage (mean 96.3%), two samples failed only because of excessive minor variant counts (>20 variations at ≥5% frequency). With a mean reference coverage of 82.1% and a median coverage depth of 937X, the failed samples overall had a substantially lower average sequencing output of 13,082 reads than the successful samples. MIRA-generated details are given in the supplementary table S1.

### 3.5. Genotype and Phylogenetic Analysis Reveals International Clusters

#### 3.5.1 Genotyping and Lineage Classification of RSV-A in Bangladesh

The ON1 genotype was found in all 43 RSV-A study sequences from Bangladesh, of which 35 are SARI cases and 8 are ILI cases. Significant variation within the local cohort was identified by subsequent lineage assignment (Figure 4). The A.D.3.7 lineage accounted for the bulk of strains (n=27) (22 SARI cases and 5 ILI cases), which clustered widely with global sequences from the USA, France, Spain, and South Africa. Within the A.D.3 clade, a clear, tight cluster of eight strains (five SARI cases and three ILI cases) exhibited phylogenetic affinity with sequences from comparable areas (USA, France, Spain, and South Africa).

**Figure 4.**
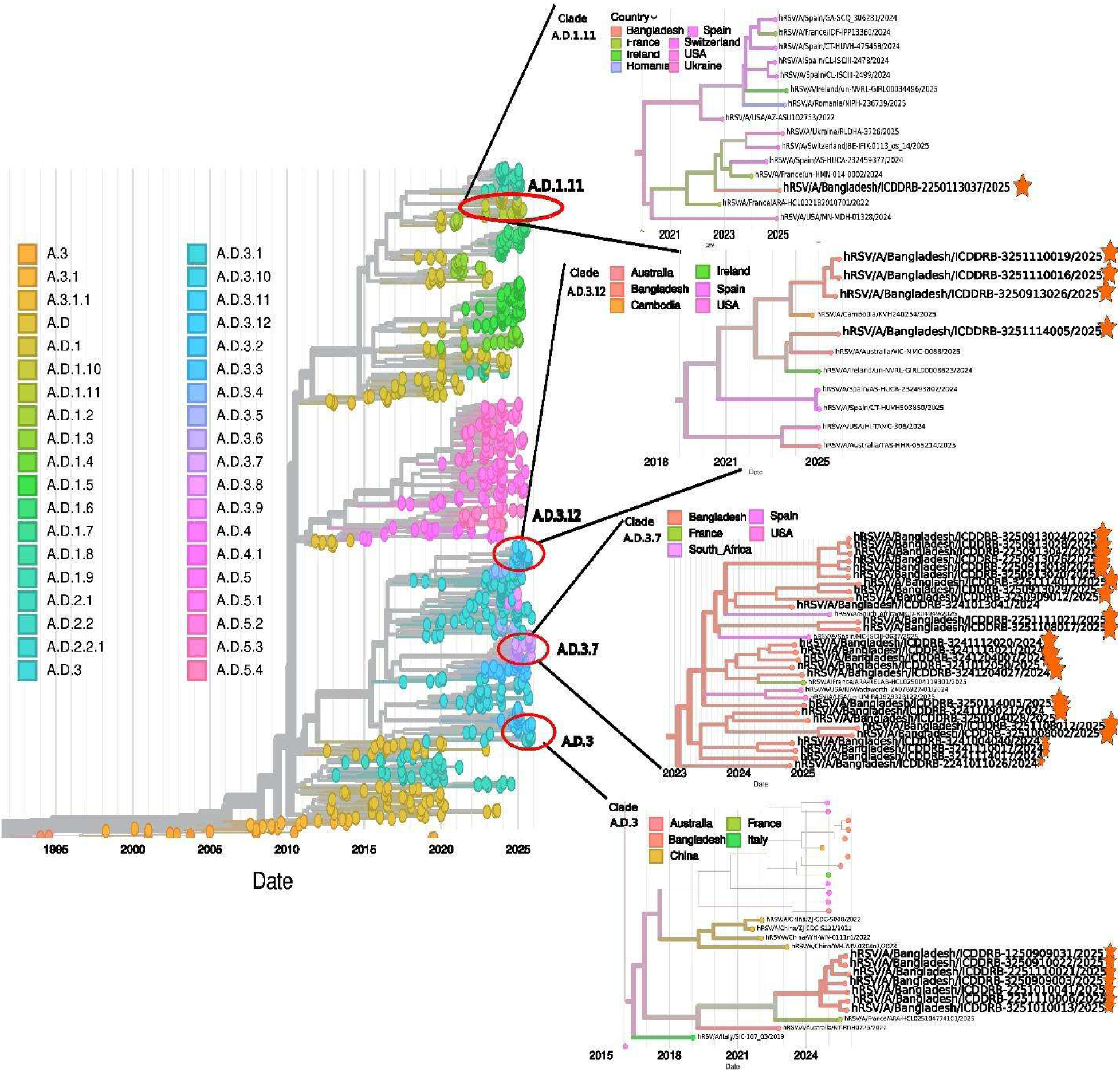
Time-resolved whole-genome phylogeny of RSV-A sequences from Bangladesh (2024–2025) generated using Nextstrain/Augur. Bangladeshi sequences (highlighted) cluster within the ON1 genotype across five sub-lineages: A.D.3.7 (n=27, 63%), A.D.3 (n=8, 19%), A.D.3.12 (n=4, 9%), A.D.3.1 (n=3, 7%), and A.D.1.11 (n=1, 2%), each phylogenetically interspersed with global reference sequences from North America, Europe, Southern Africa, and Oceania, indicating multiple independent introductions into Bangladesh.

Furthermore, four strains (all SARI cases) that merged with isolates from the USA, France, Spain, and South Africa were assigned to the A.D.3.12 lineage. The Bangladeshi sequences form a distinct cluster within the A.D.3.1 lineage, probably reflecting a localized transmission chain that diverged from the lineages circulating in the USA, France, or Oceania, such as Australia and New Zealand. Three strains (all SARI cases) formed a tight cluster within the A.D.3.1 lineage. The A.D.1.11 lineage was finally ascribed to a single strain (the SARI case), which clustered with international sequences from Spain, Switzerland, France, the USA, Ukraine, and Romania. The co-circulation of these several lineages points to several separate RSV-A variant introduction events into Bangladesh.

#### 3.5.2. Genotyping and Lineage Classification of RSV-B in Bangladesh

It was determined that all six RSV-B study sequences from Bangladesh—all of which are SARI cases—belong to the BA9 genotype. The majority of these strains (n = 5) were assigned to the B.D.E.1 lineage by phylogenetic lineage assignment; these sequences closely clustered with strains that have recently been reported from the USA and other European nations (Figure 5). The remaining strain was placed in the B.D.E.1.2 sub-lineage, a clade linked to recent North American outbreaks.

**Figure 5.**
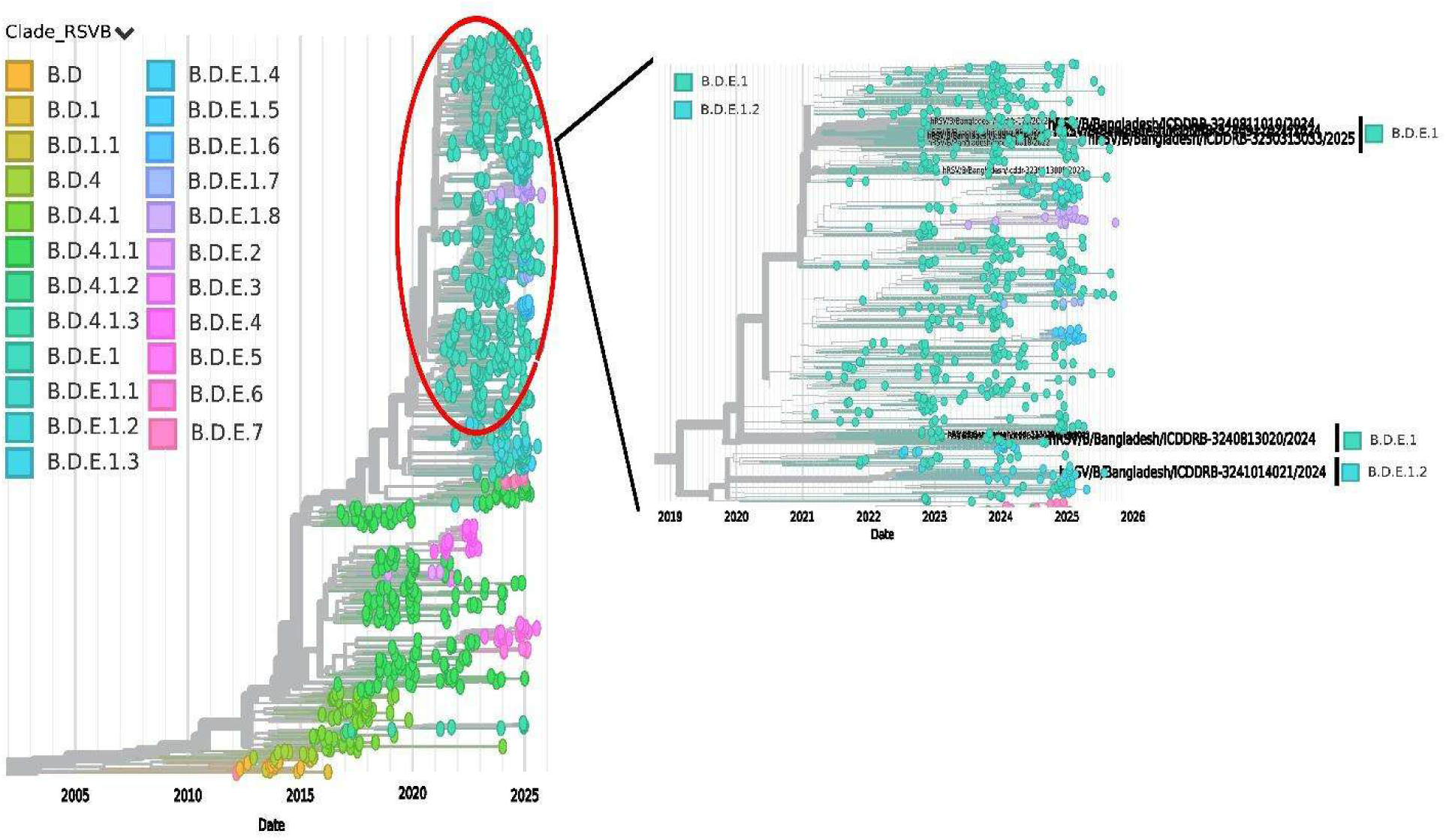
Phylogenetic study of Bangladeshi RSV-B sequences. Six locally produced strains from the BA9 genotype are included in the phylogenetic tree and are indicated in green next to global reference sequences. One set of these strains clusters within the B.D.E.1.2 sub-lineage (bold), while five strains cluster within the B.D.E.1 lineage (bold).

### 3.6. Overall Mutation

#### 3.6.1. Amino Acid Mutation Burden Across the hRSV-A Genome

Whole-genome sequencing of 43 hRSV-A cases from Bangladesh (2024–2025) identified 1,796 amino acid substitutions (187 unique; per-sequence burden 33–52, mean 41.8 ± 4.2). A significantly uneven distribution of mutational load throughout the genome was revealed by gene-wise analysis (Figure 6). Eleven mutations were fully fixed (100%) across all sequences—five in G (H90Y, I134K, G224E, I265L, D284G), two in L (R256K, Y598H), and one each in M2-1 (S176P), M2-2 (Y24C), N (V352A), and P (L55P)—defining a robust lineage-specific backbone. Six mutations (H67N, G106R, R151Q, K221R, P274L, and T319K) co-occurred in exactly 27/43 sequences (62.8%), and two outlier strains (ICDDRB-3241114017/2024, ICDDRB-3241109021/2024) shared an identical 21 substitutions and more than 60 total genome-wide mutations (Figure 6).

**Figure 6:**
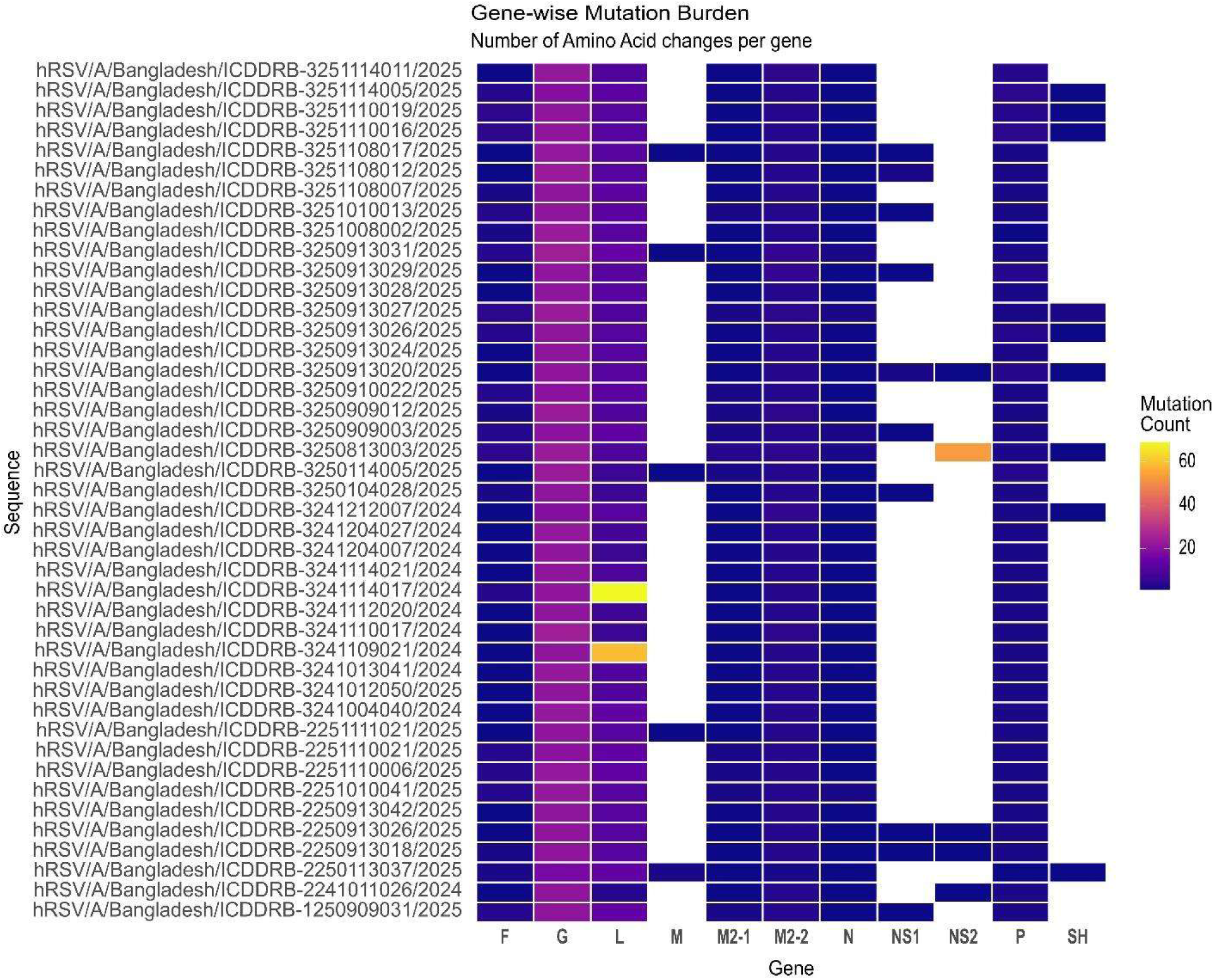
Gene-wise mutation burden summary in RSVA

With a C-terminal haplotype block (N1723S, E1725G, G1731D) in 32/43 (74.4%) and R2117K in 27/43 (62.8%), the L gene came in second (429 substitutions, 51 unique locations; mean 10.0 ± 2.8), suggesting related evolutionary alterations in the polymerase C-terminal domain. Crucially, the F gene contained 83 alterations spread over 16 locations; T12I was very universal (42/43, 97.7%), while S276N, which is found in antigenic site II and is targeted by nirsevimab and palivizumab, was present in 15/43 (34.9%), necessitating prompt phenotypic observation for possible antibody escape. Three fixed/near-fixed mutations (Y24C, S44N, and T77A) and six uncommon changes observed only once were among the 139 substitutions (11 unique locations) found in M2-2.

The N and M2-1 genes were very stable, with only one fixed change each (V352A and S176P). Finally, 89 out of 187 unique changes (47.6%) were found in only one virus each (private mutations), mostly in the L gene (29), G gene (26), F gene (9), and M2-2 gene (6). A complete catalogue of all gene-wise mutations with their positional coordinates, sequence frequencies, and prevalence classifications is provided in the supplementary tables S2 to S12.

#### 3.6.2. Amino Acid Mutation Burden Across the hRSV-B Genome

Across the hRSV-B Genome, we analyzed six hRSV-B genomes from Bangladesh (2024–2025) and found 173 amino acid changes (75 unique types). Each virus had an average of 28.8 modifications per sequence, ranging from 26 to 32 (Figure 7). In contrast to RSV-A, only three changes—F: S190N, G: S100G, and M2-2: I2T—were detected in five of the six viruses (83%). With 105 mutations spread across 29 locations (an average of 17.5 per virus), the G gene was once more the most varied. Four of the six viruses shared a set of seven G gene alterations (I137T, P214S, P221L, I252T, I268T, S275P, and Y285H), which combined formed a distinctive G gene pattern.

**Figure 7:**
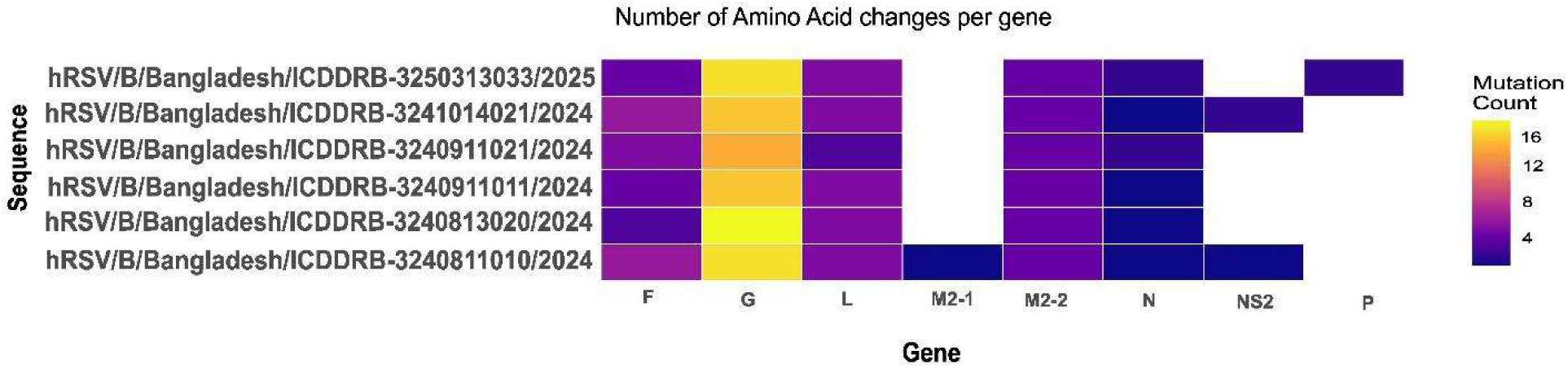
Gene-wise mutation burden summary in RSVB

The L gene had 40 changes (average 6.7 per virus), with K570R and R1759K being the most common. The F gene had 12 changes, but none involved the antibody target site (antigenic site II), meaning resistance to palivizumab or nirsevimab was not seen. The M2-2 gene carried a four-change block (I2T, M27T, D35N, C49F) in 4 of 6 viruses. Most notably, 48 of 75 unique changes (64%) were found in only one virus each -so-called private mutations - spread across G (19), L (13), F (5), and M2-2 (4). A complete catalogue of all gene-wise mutations with their positional coordinates, sequence frequencies, and prevalence classifications is provided in the supplementary tables **S13** to **S20.**

#### 3.6.3. Mutational Landscape of the RSV Fusion (F) Protein in Bangladesh

Analysis of the F protein across 43 RSV-A and 6 RSV-B sequences from recent Bangladeshi isolates (2024–2025) showed a remarkably asymmetric mutational landscape between the two subtypes (Figure 8). The mutational profile in RSV-A was dominated by the near-ubiquitous T12I substitution (97.7%), which is found inside the signal peptide. This location has previously been identified as one of the most changeable non-antigenic areas of the F gene [48].

**Figure 8.**
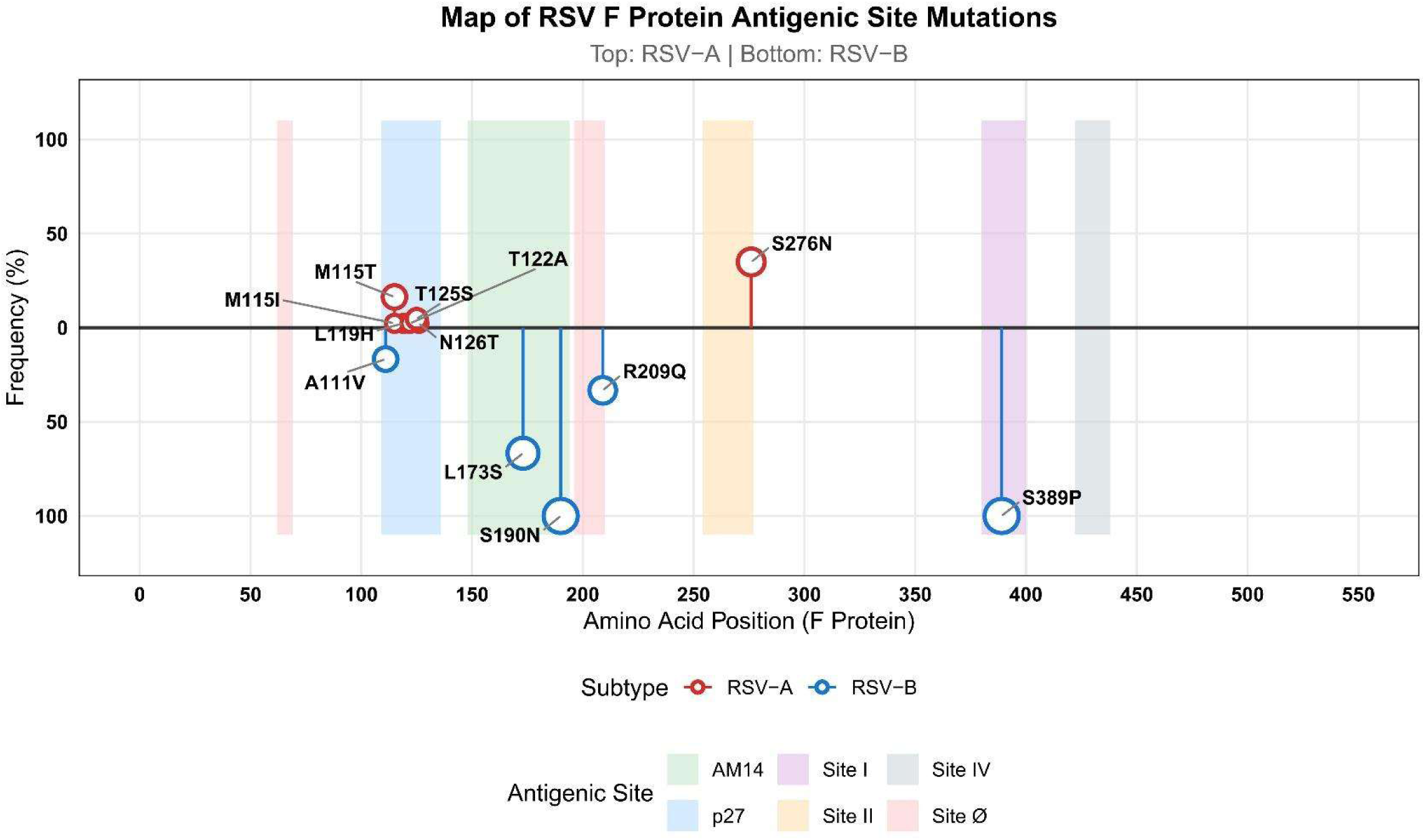
Map of RSV F Protein Antigenic Site Mutations in Contemporary Bangladeshi Isolates (2024–2025). The frequency (%) of amino acid changes in the RSV fusion (F) protein’s antigenic sites, found in 43 RSV-A (top, red) and 6 RSV-B (bottom, blue) sequences from Bangladesh, is shown in the lollipop plot. The y-axis shows the frequency of mutations as a percentage of sequences carrying each substitution, while the x-axis shows the amino acid position along the F protein (1–574). Every lollipop stem stretches from the zero baseline to the circle, the size of which is correlated with the frequency of mutations. The boundaries of recognized antigenic sites are indicated by colored vertical bands: Site Ø (pink), p27 (blue), AM14/α2α3β3β4 (green), Site II (yellow), Site I (purple), and Site IV (gray).

Furthermore, antigenic site II contained S276N in 34.9% of RSV-A sequences. This mutation, which maps directly to the palivizumab-adjacent region of the F protein and has been reported in modern genotypes at frequencies up to 70–99% worldwide, raises concerns about changing antibody recognition at this therapeutically significant site. M115T (16.3%), T125S (4.7%), L119H, T122A, and N126T were additional RSV-A variations in the p27 antigenic location that clustered inside the F protein’s most variable pre-fusion antigenic area.

RSV-B sequences, on the other hand, showed a much more cohesive mutational signature: L173S was found in 66.7% of sequences, R209Q in 33.3%, and S190N, S211N, and S389P each appeared at 100% frequency. A significant, subtype-defining divergence from the historical RSV/A Long reference strain used in the majority of current vaccine formulations is highlighted by the fixation of S389P at antigenic site I in all RSV-B isolates, which was previously reported exclusively in RSV-B at 100% frequency and absent from RSV-A. Together, these results demonstrate that modern RSV circulating in Bangladesh carries mutations at immunologically relevant F protein sites. For RSV-B, the fixation of S389P at antigenic site I and the S190N/S211N substitutions represent the established mutational signature of B.D.E lineages. Crucially, none of these mutations are associated with resistance to nirsevimab, palivizumab, or current maternal vaccines, though continued monitoring of these positions as countermeasure deployment scales is warranted

#### 3.6.4. Mutational Landscape of the RSV Attachment (G) Protein in Bangladesh

The G protein of Bangladeshi isolates of RSV-A (ON1 genotype, n=43) and RSV-B (BA9 genotype, n=6) showed significant amino acid variability concentrated across the hypervariable areas, yet the two subtypes’ mutational fingerprints were remarkably different.

In RSV-A, the mutational burden was dominated by near-universal substitutions in both HVR1 and HVR2 — H90Y, I134K, and T113I reached 100%, 100%, and 97.7% frequency, respectively, in HVR1, while HVR2 harbored equally fixed changes, including G224E, I265L, and D284G at 100%, alongside H258Q and H266L at 97.7%. The central conserved domain (CCD), which is traditionally thought to be immunologically stable, was not spared; N178G, a substitution found in the heparin-binding domain of 97.7% of RSV-A sequences, may have an impact on CX3CR1-mediated host cell attachment, a mechanism that is becoming more and more important to RSV pathogenesis. In contrast, RSV-B showed a more streamlined but equally fixed mutational repertoire: A74V, S100G, T131A, and I137T were all fixed at 100% frequency in HVR1, and HVR2 mutations such as P214S, P221L, I252T, I268T, S275P, and Y285H also reached 100% frequency, indicating a nearly total divergence from the RSV-A reference backbone across almost every structural domain of the G protein (Figure 9). Significantly, 16.7% of RSV-B sequences included a premature stop codon (*311Q), which raises concerns for shortened G protein production and its possible consequences for immune evasion. Together, these results highlight the antigenic gap between circulating strains and historical reference sequences employed in vaccine development by revealing that modern Bangladeshi RSV strains possess highly fixed, subtype-defining G protein mutations encompassing both variable and conserved domains.

**Figure 9:**
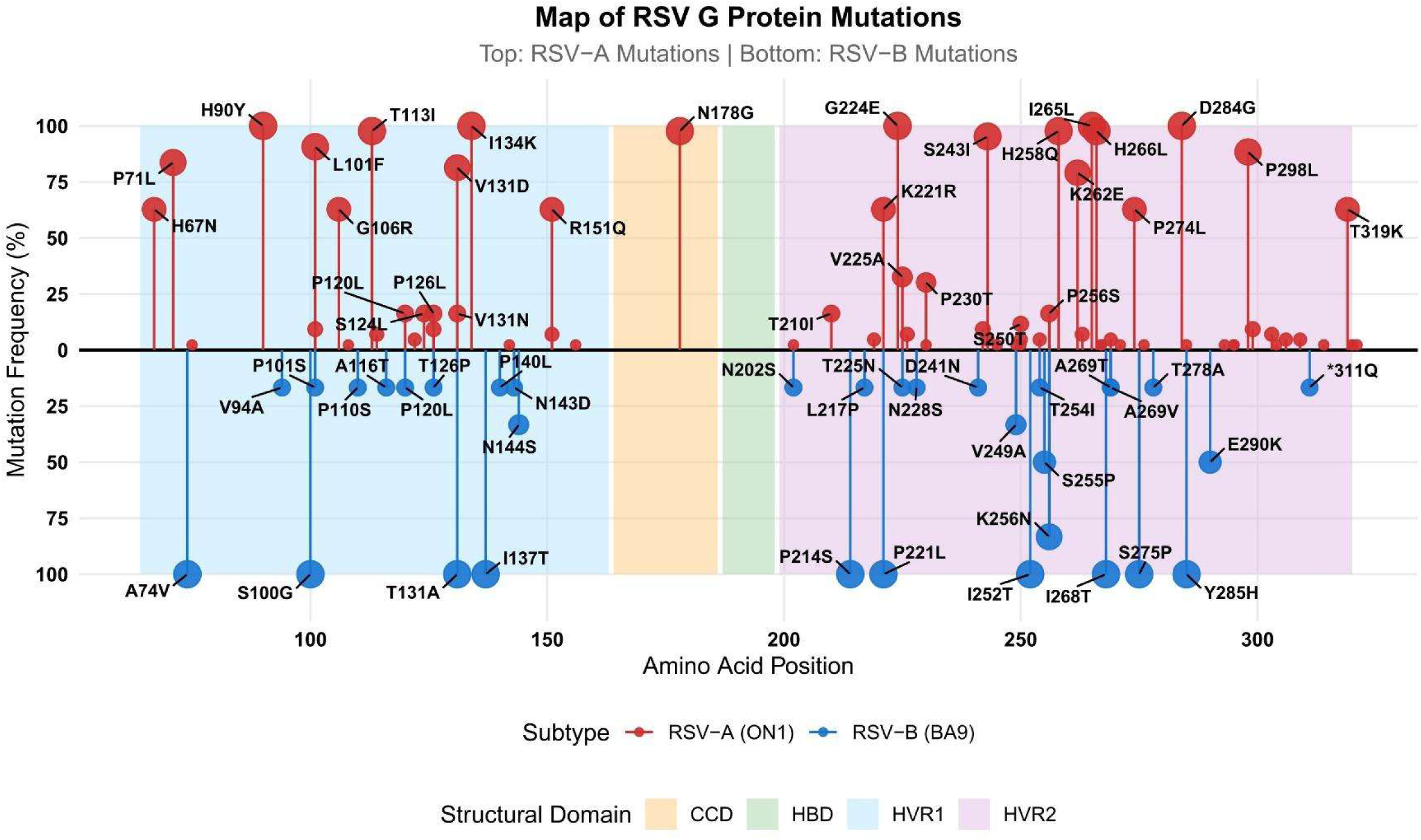
Map of RSV G Protein Amino Acid Substitutions in Contemporary Bangladeshi Isolates (2024–2025).

The frequency (%) of amino acid alterations in the RSV attachment (G) protein found in 43 RSV-A (ON1 genotype, top, red) and 6 RSV-B (BA9 genotype, bottom, blue) sequences from Bangladesh are shown in the lollipop plot. The y-axis shows the replacement frequency as a percentage of sequences containing each alteration, while the x-axis shows the location of amino acids along the G protein.

Each lollipop stem extends from the zero baseline to the circle, with circle size proportional to substitution frequency. Only substitutions with frequency greater than 10% are labeled for clarity, as defined in the analysis code. Colored background bands denote the boundaries of known structural domains: HVR1 — first hypervariable region (blue, aa 64–163), CCD — central conserved domain (orange, aa 164–186), HBD — heparin-binding domain (green, aa 187–198), and HVR2 — second hypervariable region (purple, aa 199–320).

#### 3.6.5. Predicted N-Glycosylation Landscape of the RSV

##### 3.6.5.1. N-linked Glycosylation Profiling of the RSV Fusion Protein in Bangladeshi Strains

N-linked glycosylation of the RSV fusion (F) protein is a key determinant of immune evasion, as glycan shields restrict antibody access to conserved neutralizing epitopes[19, 20].To characterize the glycosylation landscape of circulating Bangladeshi RSV strains (2024–2025), we predicted N-glycosylation sequons (N-X-S/T) in the F proteins of 43 RSV-A and 6 RSV-B isolates using NetNGlyc 1.0 (threshold ≥ 0.5)[21]. The ancestral RSV-A reference (NC_001803.1) carried seven co nfirmed sites at positions N232, N252, N264, N338, N353, N456, and N489. Strikingly, all 43 Bangladeshi RSV-A isolates uniformly acquired a novel, high-confidence glycosylation site at position N75 (sequon NKTR; jury score 9/9; potential ∼0.79) — absent in the reference — representing a newly fixed glycan addition in circulating strains (Figure 10a). Moreover, most isolates displayed a duplication of the C-terminal glycosylation region, carrying two closely spaced confirmed sites at N487 and N490 (both NYSN sequons) instead of the single reference N489, effectively doubling local glycan density. Additionally, a significant portion gained a site at N453–N456, an immunologically accessible surface region, indicating active glycan modification close to important antigenic zones. The universal fixation of N75 across all Bangladeshi isolates has not been explicitly reported in global datasets, highlighting the epidemiological significance of this observation. These findings are consistent with global surveillance, where gains and losses of F protein glycosylation sites in circulating RSV-A strains have been linked to altered antigenic presentation and potential immune escape[2, 22]. In sharp contrast, RSV-B F proteins from all six Bangladeshi isolates shared an identical set of five conserved glycosylation sites (N27, N70, N116, N120, N126) with near-identical prediction scores (average ∼0.742), mirroring global reports of strong purifying selection on the RSV-B F protein (Figure 10b). [23]. Collectively, these findings reveal a subtype-divergent glycan remodeling strategy — RSV-A undergoing progressive glycan expansion relative to the reference, while RSV-B maintains strict glycan conservation — with important implications for the efficacy of current pre-fusion F-targeting vaccines (Arexvy (GSK), Abrysvo (Pfizer), and mRESVIA (Moderna)) and monoclonal antibodies such as nirsevimab.[19, 24]. A complete catalogue of all N-linked Glycosylation Profiling of the RSV Fusion Protein in Bangladesh is provided in the supplementary tables S21 to S25.

**Figure 10a.**
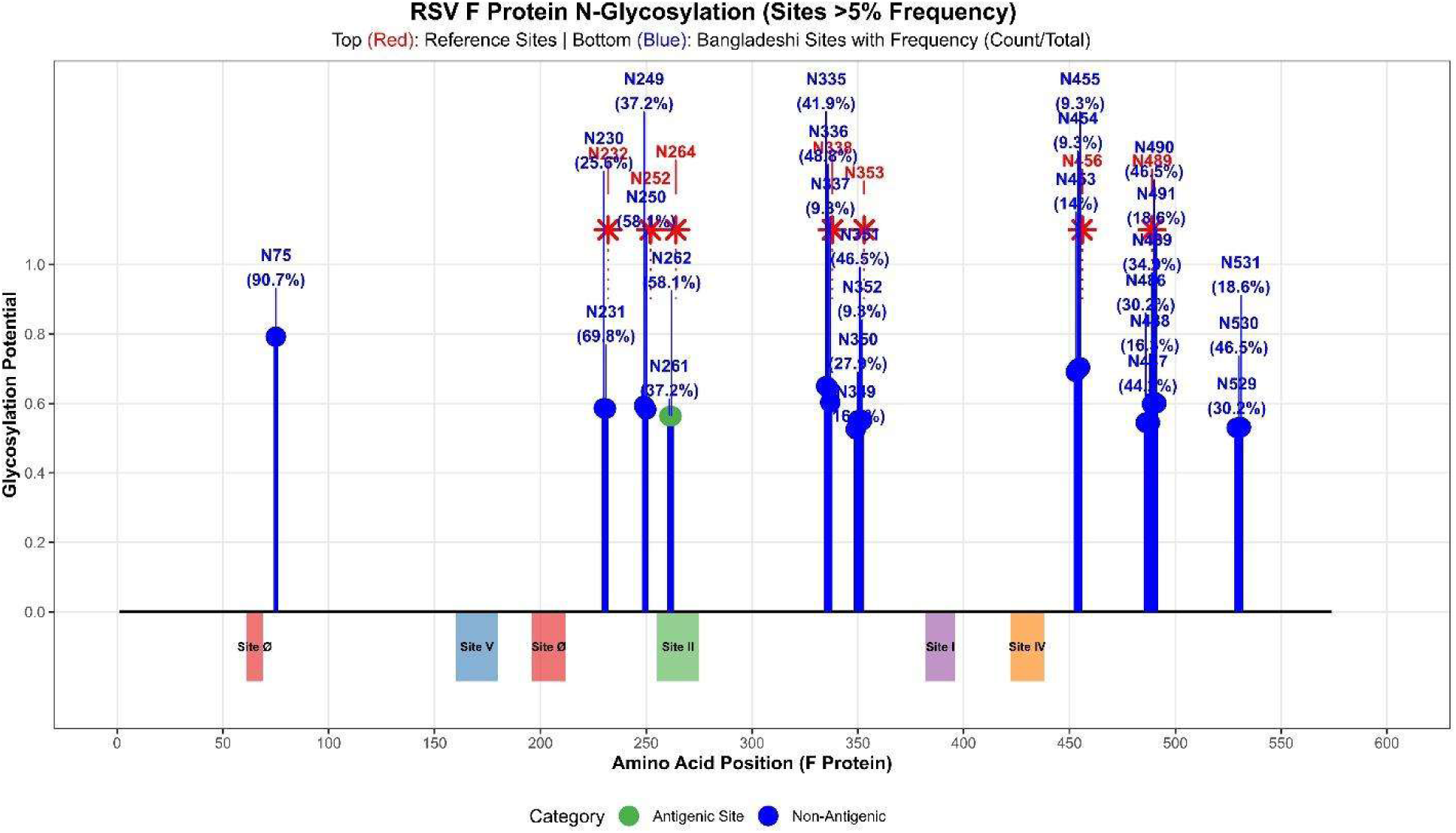
Comparative N-glycosylation landscape of the RSV-A F protein in Bangladeshi circulating strains (2024–2025) versus the reference genome (NC_001803.1). Blue stems indicate glycosylation sites predicted in circulating Bangladeshi isolates (n=34); red stars indicate reference sites. All 34 isolates uniformly carry a new N-glycosylation site at N75 (absent from the reference), and show duplication of the C-terminal glycan region (N487+N490 replacing single reference N489), representing progressive glycan expansion compared to the reference backbone.

**Figure 10b.**
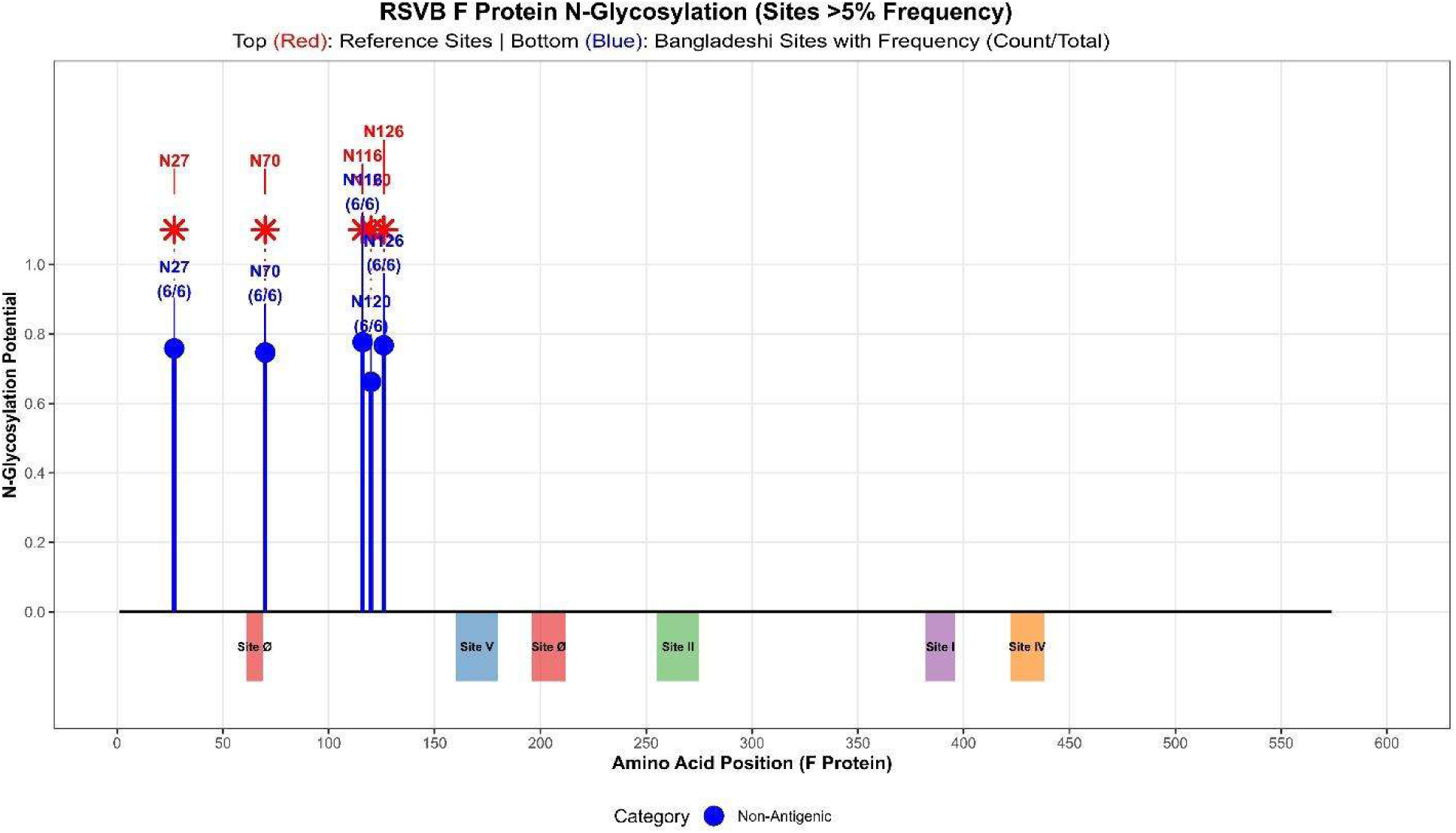
Comparative N-glycosylation landscape of the RSV-B F protein in Bangladeshi circulating strains versus the reference genome. RSV-B F proteins from all six Bangladeshi isolates conserve an identical set of five canonical glycosylation sites (N27, N70, N116, N120, N126), mirroring the reference and consistent with strong purifying selection on the RSV-B F glycan scaffold.

##### 3.6.5.2. N-Glycosylation Landscape of the RSV G Protein in Bangladeshi Strains

N-glycosylation profiling of the RSV G protein using NetNGlyc 1.0 (score threshold >0.5) revealed biologically significant and subgroup-distinct glycosylation architectures in Bangladeshi circulating strains. For RSV-A, the reference genome (NC_001803) harbors six predicted N-glycosylation sequons - N5 in the cytoplasmic tail (predicted but biologically inaccessible to ER-mediated glycosylation machinery) [25], N108 in mucin-like domain I [26] and N242, N255, N256, and N323 within the second mucin-like domain (HVR2) [27, 28] - the most antigenically variable region of the G protein (Figure 11a). Bangladeshi RSV-A strains preserved this multi-site scaffold, consistent with prior Bangladesh surveillance identifying equivalent HVR2 glycosylation sequons [8] and with the established global pattern of RSV-A carrying more N-glycosylation sites than RSV-B, conferring broader epitope masking [29].

**Figure 11a.**
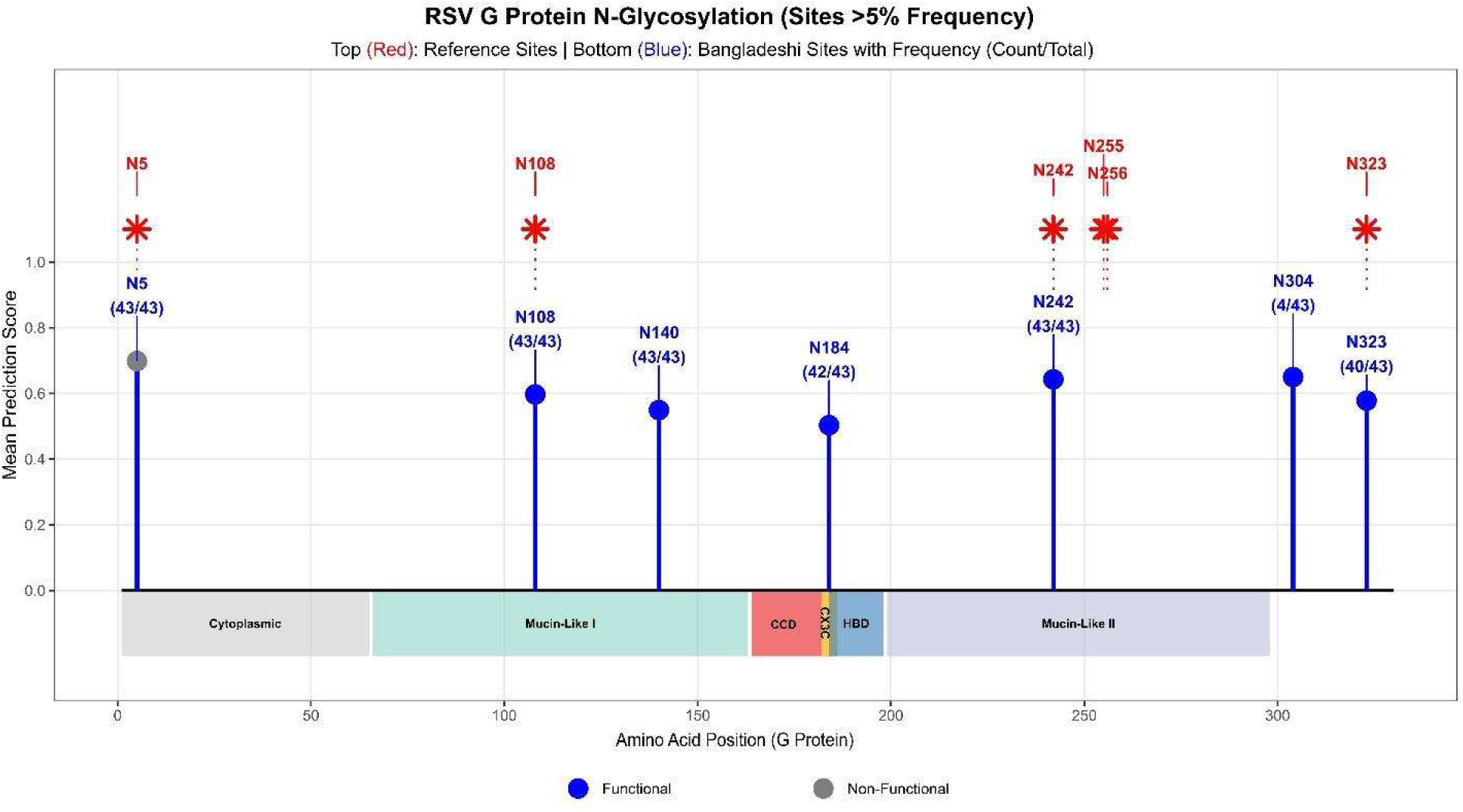
Comparative N-glycosylation landscape of the RSV-A G protein (n=43). Bangladeshi isolates preserve the reference multi-site HVR2 scaffold; no novel acquisitions or losses were detected in the HVR2 sequons (N242, N255, N256, N323), consistent with prior Bangladesh surveillance.

Three conserved sites for RSV-B, N86, N228, and N294, are present in the Australian reference strain (OP975389.1) and are representative of the BA lineage core that has been documented worldwide [30, 31]. These sites are universally retained across all six Bangladeshi RSV-B sequences from 2024–2025, highlighting the strong evolutionary constraint on these positions (Figure 11b). Importantly, D253N/K258N glycan acquisitions at equivalent HVR2 positions in European strains [30] and S100N-driven glycan gain in post-pandemic RSV-A in the US (Rios-Guzman et al., 2024) were mirrored by 67% (4/6) of Bangladeshi RSV-B strains gaining an additional site at N256 within mucin-like domain II.

**Figure 11b.**
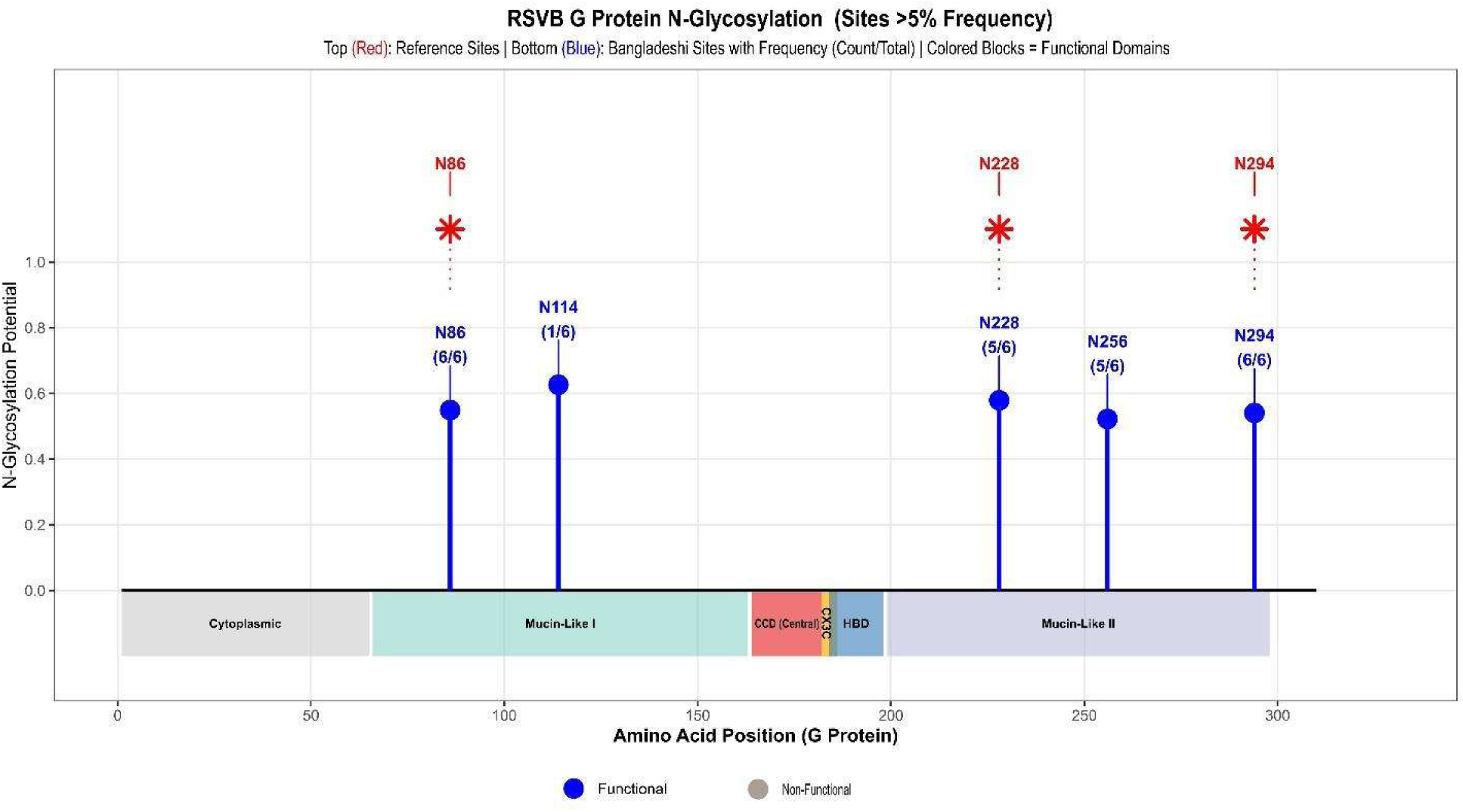
Comparative N-glycosylation landscape of the RSV-B G protein (n=6). All six Bangladeshi RSV-B isolates retain three conserved core glycosylation sites (N86, N228, N294). Notably, 67% (4/6) acquired an additional HVR2 site at N256, paralleling glycan acquisitions reported in European B.D.E lineages and suggesting convergent evolutionary pressure toward a denser G protein glycan shield.

These modifications physically block antibody access to important neutralizing epitopes, which is a known RSV immune evasion strategy [32]. This has obvious consequences for treatments that target G proteins. Collectively, Bangladeshi RSV strains are actively evolving toward a denser N-glycan shield, a convergent signal across Asia, Europe, and North America, warranting systematic glycan-aware genomic surveillance. A complete catalogue of all N-linked Glycosylation Profiling of the RSV G Protein in Bangladesh is provided in the supplementary tables S26 to S27.

#### 3.6.6. Predicted O-Glycosylation Landscape of the RSV

##### 3.6.6.1. O-Glycosylation Landscape of the RSV F Protein in Bangladeshi Strains

O-linked glycosylation profiling of the RSV fusion (F) protein across both RSV-A and RSV-B subgroups uncovered a rich, subgroup-differentiated landscape of mucin-type modifications at serine and threonine residues spanning the full ectodomain. For RSV-A, predicted O-glycosylation occupancy varied substantially between globally representative reference sequences and locally circulating BD strains — with certain positions in the F2 subunit and near the p27 cleavage region showing increased frequency in BD strains (e.g., T67, T140, T173: +0.03 to +0.06), while others, particularly around antigenic site II (S275: −0.06) and the p27 region (S107: −0.05), were selectively depleted — indicating active, immune-pressure-driven remodeling of the glycan landscape in circulating RSV-A (Figure 12a) [29, 33]. For RSV-B, O-glycosylation was both more abundant and more conserved, with a clearly stratified frequency distribution: very high-frequency sites (>85% occupancy; T67, T173/S173, T476) anchored in the F2 subunit and at antigenic site Ø and site V — the target regions of nirsevimab and suptavumab, respectively — alongside moderate-frequency sites (50–75%) distributed across F1 (Figure 12b) [34, 35]. Critically, in both subgroups, the highest-occupancy O-glycosylation sites co-localized with or directly flanked key therapeutic antibody epitopes, including site II (T262, S255; palivizumab target), site IV (T390; MK-1654 target), and site Ø (T476; nirsevimab target), raising the important possibility that O-glycan additions at these positions act synergistically with amino acid variation to reduce antibody accessibility — a mechanism globally analogous to the glycan shielding described for influenza hemagglutinin and SARS-CoV-2 spike [36]. RSV-B exhibited a notably higher mean O-glycan site frequency and a greater proportion of very high-frequency sites than RSV-A, consistent with its broader antigenic divergence and higher overall F protein variability [28, 37]. Together, these data represent a frequency-resolved, residue-level O-glycan map of the RSV F protein for both subgroups, positioning O-glycosylation as a dynamic, selection-responsive arm of RSV immune evasion that differs meaningfully between RSV-A and RSV-B and warrants explicit consideration in the rational design of broadly protective, subgroup-inclusive RSV vaccines and therapeutics [24, 29]. A complete catalogue of all O-linked Glycosylation Profiling of the RSV F Protein in Bangladesh is provided in the supplementary tables S28 to S29.

**Figure 12a:**
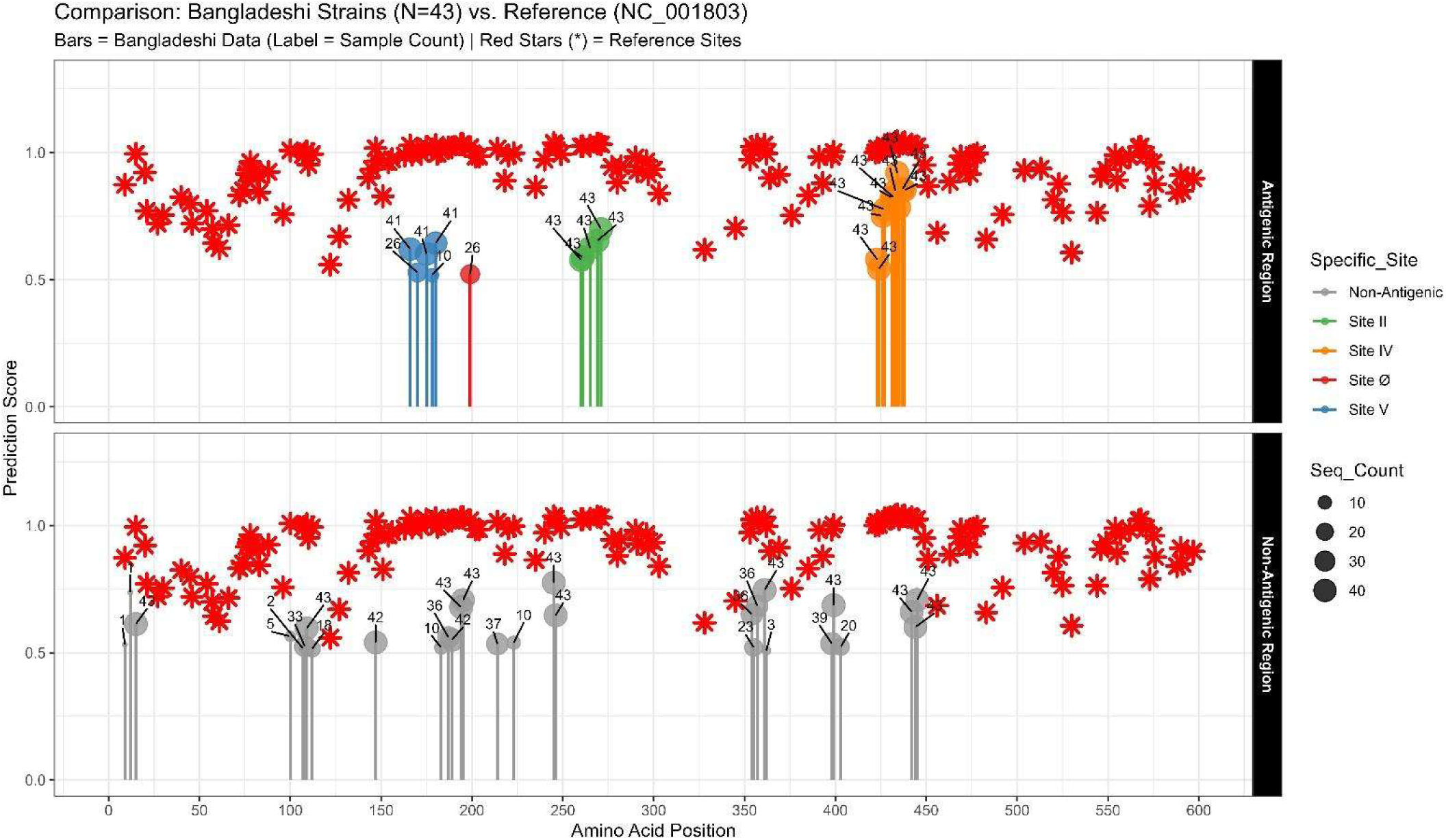
O-glycosylation site occupancy comparison between RSV-A reference and BD circulating strains across the F protein. Comparative lollipop or bar plot displaying predicted O-linked glycosylation frequency (proportion of sequences, NetOGlyc 4.0, G-score ≥ 0.5) at serine (S) and threonine (T) residues across the RSV-A F protein ectodomain for globally representative reference sequences (grey) and locally circulating BD strains (colored). Residue positions with differential occupancy (|Δ| > 0.04) are labeled. Dashed vertical lines demarcate F2, p27, and F1 subunit boundaries; horizontal dashed lines indicate antigenic site regions (Ø, II, IV, V).

**Figure 12b:**
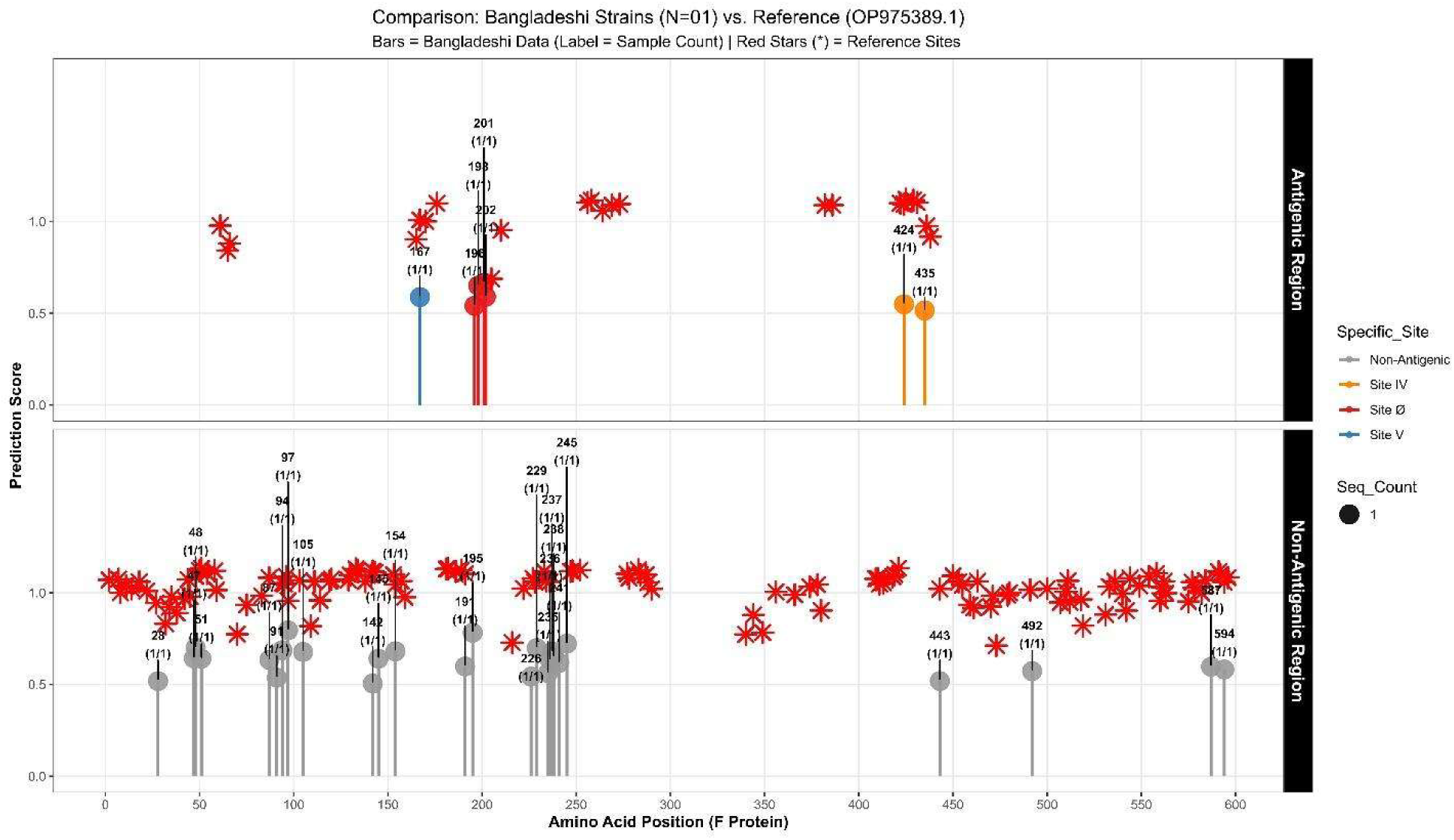
Frequency-resolved O-glycosylation map of the RSV-B F protein across circulating strains. Lollipop plot illustrating predicted O-linked glycosylation frequency (NetOGlyc 4.0, G-score ≥ 0.5) at individual serine and threonine residues across the RSV-B F protein ectodomain. Occupancy frequency is represented by stem height (0–1.0). Sites with the highest occupancy or closest antigen are shown by labeled residues. The F2 (blue), p27 (orange), and F1 (green) sections are distinguished by color coding. The borders of important antigenic sites (Ø, I, II, IV, and V) are indicated by dashed lines. Very high-frequency sites (≥ 85%) are highlighted.

##### 3.6.6.2 O-Glycosylation Landscape of the RSV G Protein in Bangladeshi Strains

Using NetOGlyc 4.0, O-glycosylation of the RO-glycosylation of the RSV attachment (G) protein was predicted for 43 RSV-A and 6 RSV-B clinical isolates that were gathered in Dhaka, Bangladesh (2024–2025). The mucin-like domain I (MLR-I; aa 66–163) and the central conserved domain (CCD; aa 164–186) of the G protein contain the majority of the 29 high-confidence O-glycosylation sites (NetOGlyc score >0.5) for RSV-A, which cover positions 81–182. Sites at positions 85, 86, 88, 89, 112, 113, 117, 119, 120, 158, and 176 were detected in all 43 isolates (100% frequency), with an additional 7 sites (positions 142–152) showing near-universal prevalence (≥95%), indicating a strikingly stable glycan landscape across the Bangladesh RSV-A population (Figure 13a). Interestingly, position 176—within the CCD—was glycosylated at 100% frequency in our isolates, which is in line with previous findings that this region is crucial for glycan-mediated shielding next to the CX3C motif (aa 182–186) [24, 38, 39]. 9.3% of RSV-A strains were projected to be glycosylated at position 182, which is part of the CX3C chemokine-mimicry motif itself. This prediction is consistent with prior findings that partial glycosylation at this site can modify CX3CR1 interaction [40]. For RSV-B, 38 O-glycosylation sites were identified across 6 isolates spanning positions 59–236. Core mucin-like I sites at positions 66, 68, 71, 72, 74, 84, 102, 103, 104, 109, and 143 exhibited 100% frequency, all shared with the reference strain (NC_001781). Within the CCD, position 167 was glycosylated universally (6/6 isolates, 100%), and position 185 was similarly universal (6/6), while positions 182 (50%) and 176 (33%) were variably glycosylated. Intriguingly, position 186—at the C-terminal edge of the CX3C motif—was glycosylated in 1/6 RSV-B isolates and was *absent from the reference* (Figure 13b), suggesting a locally emerging glycan acquisition. The heavy and functionally organized O-glycosylation we observed at MLR-I and CCD positions in both subtypes is consistent with published evidence that mucin-type glycans in these domains form a glycan shield obscuring neutralization epitopes and dampening host antibody responses [38, 41], and our findings extend this picture to circulating Bangladeshi strains for the first time, with implications for both diagnostic and vaccine antigen design. A complete catalogue of all O-linked Glycosylation Profiling of the RSV G Protein in Bangladesh is provided in the supplementary table in S30 to S31.

**Figure 13a:**
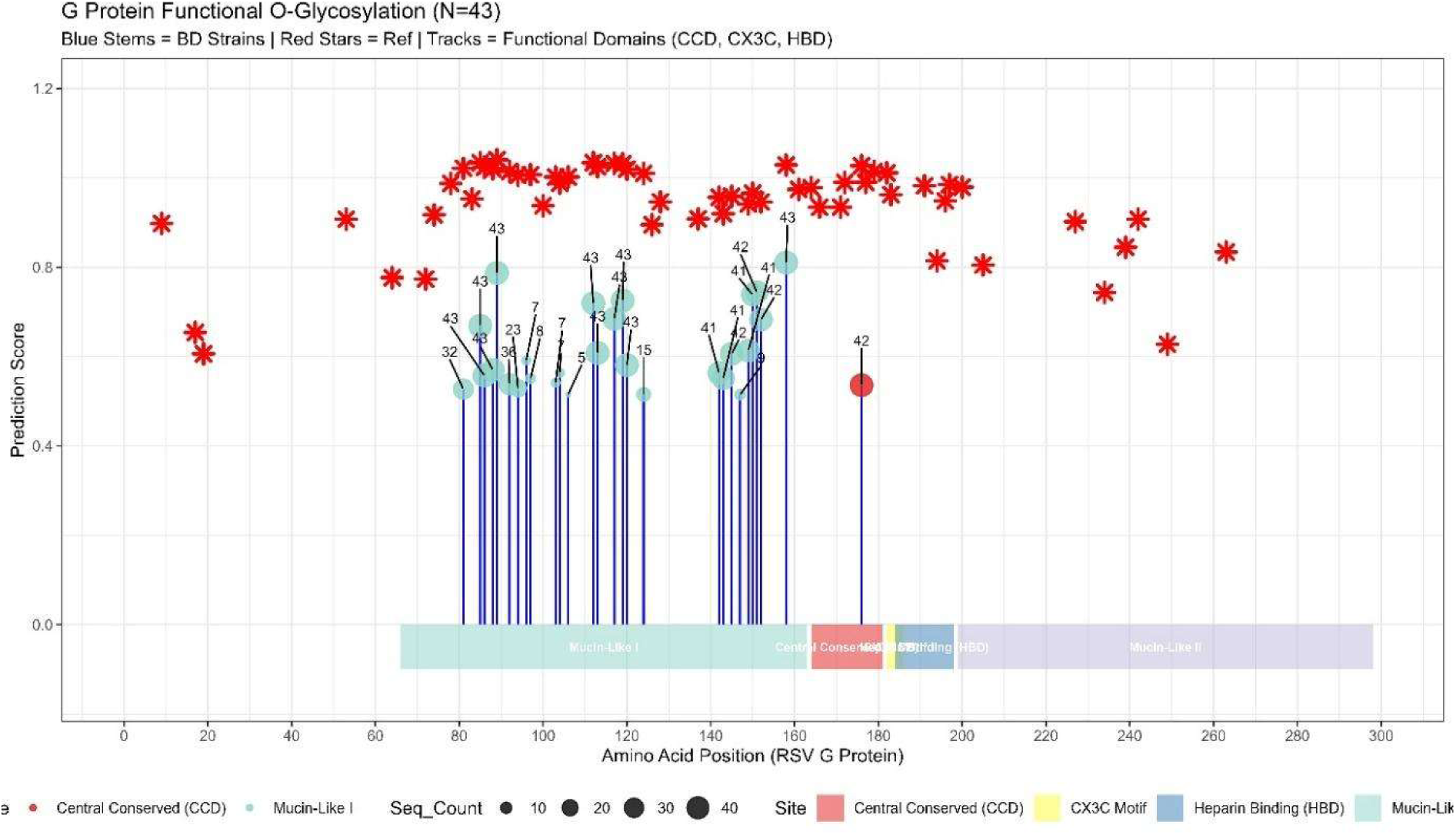
O-Glycosylation profile of the RSV-A G protein in Bangladeshi clinical isolates (N=43). Each vertical stem represents a predicted O-glycosylation site (NetOGlyc score >0.5); stem height corresponds to mean prediction score and circle size reflects the number of isolates positive at that position. Red stars indicate sites shared with the RSV-A reference strain (NC_001803). Coloured horizontal tracks denote functional domains: Mucin-Like I (teal, aa 66–163), Central Conserved Domain/CCD (red, aa 164–181), CX3C motif (yellow, aa 182–186), Heparin-Binding Domain/HBD (blue, aa 184–198), and Mucin-Like II (lavender, aa 199–298). Numbers above the stems indicate the count of positive isolates.

**Figure 13b:**
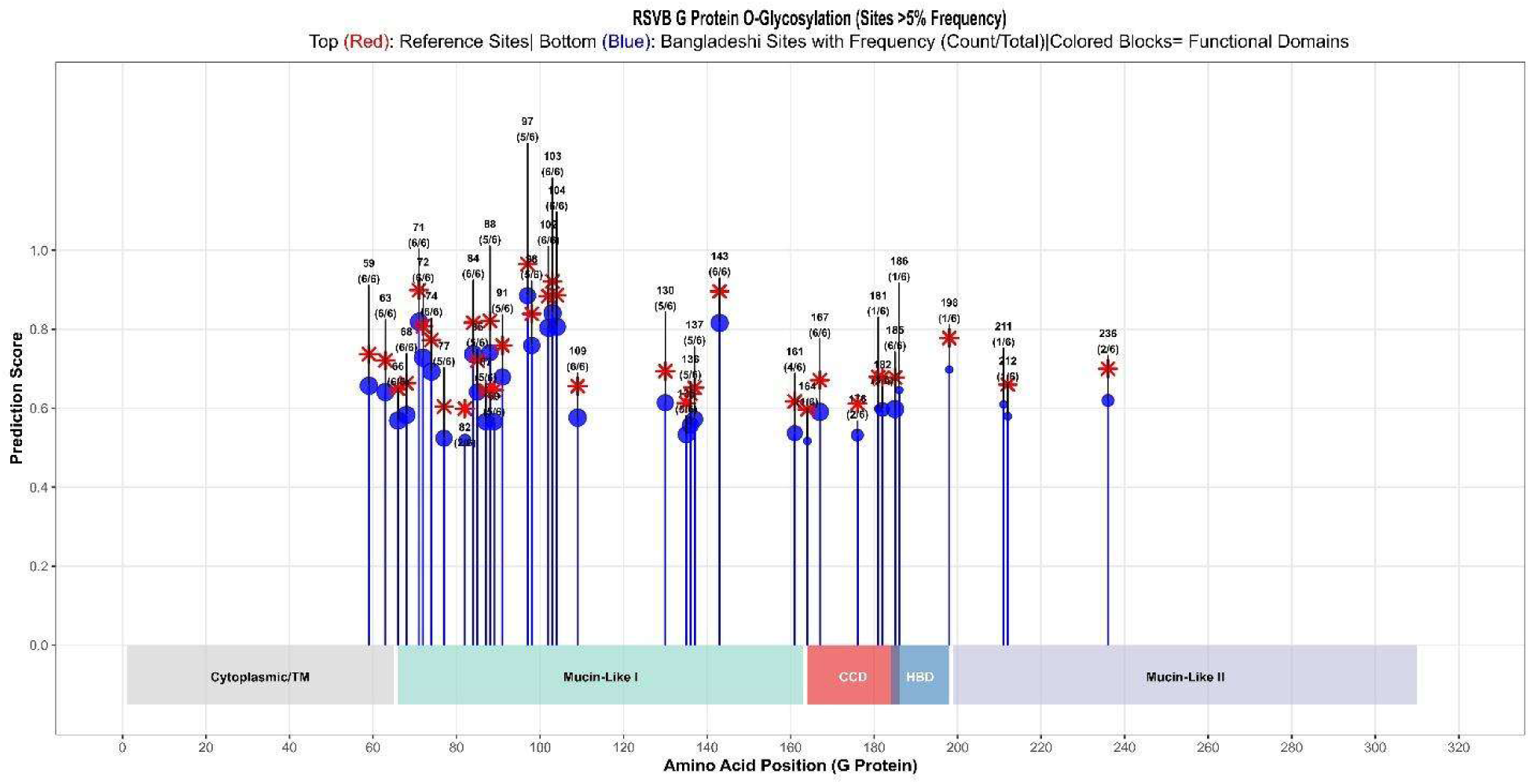
O-Glycosylation frequency profile of the RSV-B G protein in Bangladeshi clinical isolates (N=6). Stem plots depict predicted O-glycosylation sites; blue stems and circles represent Bangladeshi isolates, with point size scaled to sample count (n). Red stars mark sites shared with the RSV-B reference strain (NC_001781). Site labels show position and fraction (e.g., 6/6 = 100%). Functional domains are colour-coded as in Figure 1: Cytoplasmic/TM (grey), Mucin-Like I (teal), CCD (red), HBD (blue), and Mucin-Like II (lavender).

## 4. Discussion

Whole-genome sequencing data from RSV strains that were prevalent in the Bangladeshi population during the 2024–2025 respiratory season are presented in this work. Before the COVID-19 pandemic, RSV in Bangladesh exhibited seasonal patterns that were predicted, with previous partial-gene monitoring studies identifying ON1 and BA lineages as dominant [42]. In line with observations from other nations that indicate the post-pandemic RSV rebound was driven by pre-existing lineages that persisted in low-level circulation rather than newly emergent variants, the persistence of these same lineages in the current post-pandemic period indicates genomic continuity despite the epidemiological disruption [43].

The operational viability of whole-genome RSV sequencing within an established hospital-based surveillance infrastructure in a lower-middle-income country is confirmed by the sequencing success rate of 83.1% (49/59 eligible samples), mean genome completeness of 94.7%, and median depth of 1,500× using Oxford Nanopore Technology and the ARTIC primer scheme.

All 43 RSV-A sequences belonged to the ON1 genotype (lineage A.D) [44] with five sub-lineages co-circulating simultaneously: A.D.3.7 (n=27, 63%), while A.D.3, A.D.3.12, A.D.3.1, and A.D.1.11 comprised 19% (n = 8), 9% (n = 4), 7% (n = 3), and 2% (n = 1) of the dataset, respectively. This multi-lineage composition, which is consistent with the cross-continental dispersal of A.D.3 lineages reported in recent surveillance data from North America, Europe, and Southern Africa [45], strongly refutes a single importation event and instead shows multiple independent introductions of RSV-A into Bangladesh from geographically distinct sources.

A localized transmission chain after introduction is suggested by the tight A.D.3.1 sub-cluster, which diverges from closest global relatives in North America and Oceania. This is a post-pandemic feature seen in South Korea and Myanmar [46]. All six RSV-B sequences belonged to the BA9 genotype (B.D.E.1 or B.D.E.1.2), grouping with strains from the USA and Europe. This is in line with the worldwide dissemination of B.D.E.1, which caused RSV-B outbreaks in North America and Europe in 2022–2023 [47].

The presence of five distinct RSV-A lineages and sequences phylogenetically linked to six continents underscores the inadequacy of partial-gene sequencing for tracking RSV import diversity in Bangladesh.

Whole-genome analysis of 43 RSV-A sequences identified 1,796 amino acid substitutions (187 unique; mean 41.8 ± 4.2 per sequence), with the G gene accounting for ∼40% of all unique mutations — consistent with the well-established role of immune selection pressure on the attachment protein [48]. Eleven mutations were fully fixed across all RSV-A sequences, defining the lineage backbone for this season. A C-terminal haplotype block in the L polymerase (N1723S, E1725G, G1731D) was present in 74.4% of sequences, suggesting linked evolutionary change in the replication machinery — a feature of potential relevance to polymerase-targeting therapeutics under development. Two RSV-A outlier strains shared >60 genome-wide mutations and 21 shared G-gene substitutions, suggesting either a divergent import event or an accelerated local evolutionary trajectory. For RSV-B, the genome-wide mutation burden was lower (mean 28.8 per sequence), and 64% of unique changes were private mutations — characteristic of a small, recently introduced cluster. A premature stop codon (*311Q) was detected in one RSV-B G protein, raising questions about truncated G protein expression and its potential immune consequences, as has been noted for related RSV-B lineages [49]. The CCD substitution N178G in 97.7% of RSV-A G proteins — residing within the heparin-binding domain implicated in CX3CR1-mediated host attachment [40] illustrates how mutations in classically conserved domains may carry functional consequences that surveillance focused solely on annotated antigenic sites would overlook.

The RSV F protein is the target of all licensed RSV countermeasures, and the integrity of its key antigenic sites is a primary focus of surveillance. In RSV-A, the near-universal T12I substitution (97.7%) is a lineage-defining signal peptide variant with no demonstrated impact on antibody binding [48]. The S276N substitution at antigenic site II — present in 34.9% of RSV-A sequences — maps to the palivizumab binding region and has been documented globally, with a reported frequency of ∼5.2% in GISAID through end-2024, making the frequency observed here substantially higher than the global average and consistent with lineage-specific enrichment [50].

Crucially, the functional data that is currently available verify that S276N does not provide palivizumab resistance in vitro [51], and it has not been found to be a substitute for nirsevimab resistance [50]. Further diversification at the most structurally variable pre-fusion antigenic area is reflected in further p27-region variation (M115T 16.3%, T125S 4.7%, L119H, T122A, N126T) [52]. S190N, S211N, and S389P were fixed in all six sequences of RSV-B, whereas L173S was fixed in 66.7% of the sequences. These mutations represent the consolidated mutational signature of B.D.E lineages rather than new resistance events [50].

No mutations at nirsevimab binding site Ø with known neutralisation impact were detected in either subtype, consistent with the >99% conservation of this site across 25 critical positions reported in the OUTSMART-RSV surveillance study[53]. Taken together, these data support the continued immunological effectiveness of current F-protein-targeting interventions against circulating Bangladeshi RSV strains, while identifying S276N and the p27 cluster as sites warranting ongoing functional monitoring as countermeasure deployment scales.

Beyond amino acid replacement alone, the glycosylation profiles of the F and G proteins offer more biological context. In addition to duplication of the C-terminal glycosylation region (N487 and N490 replacing single reference N489) and an additional site at N453–N456, all 34 analyzed isolates of the RSV-A F protein uniformly acquired a high-confidence N-glycosylation site at N75, which is absent from the historical reference genome. In the context of post-pandemic worldwide surveillance, this pattern of progressive N-glycan expansion in circulating RSV-A F protein has been documented [54–56]. The particular N75 fixation seen in Bangladesh calls for structural characterization to evaluate its impact on antigenic surface accessibility. In contrast, RSV-B F proteins showed strict conservation of five canonical N-glycosylation sites (N27, N70, N116, N120, N126), consistent with strong purifying selection on the RSV-B F glycan scaffold [52]

Dense O-glycosylation at mucin-like domain I and CCD positions in both subgroups of the G protein, with 100% frequency at multiple sites in RSV-A (positions 85, 86, 88, 176) and RSV-B (positions 66, 68, 167), is consistent with the known function of these modifications in creating an immunological shield that limits antibody access to the G protein ectodomain [38, 41]. Similar HVR2 glycan increases observed in European B.D.E lineages [30] are paralleled by the acquisition of an N256 N-glycosylation site in 67% of RSV-B G proteins, indicating convergent evolutionary push toward a denser G protein glycan shield. Validated computer prediction algorithms provide the basis of all glycosylation discoveries. (NetNGlyc 1.0; NetOGlyc 4.0) [15] and necessitate mass spectrometry experimental confirmation, which is a logical next step for this dataset.

The results directly affect vaccination and RSV surveillance practices. An empirical base for continuing policy discussions about maternal RSV vaccination and newborn nirsevimab prophylaxis in Bangladesh is provided by the confirmation that all circulating lineages retain the important antigenic sites targeted by nirsevimab and the approved vaccines (mRESVIA, Arexvy, Abrysvo) [15]. The N75 glycan increase, which is present in all Bangladeshi RSV-A F proteins, should be included in national genomic surveillance annotations since it could be a helpful molecular marker for tracing local lineages.

A region that has long been underrepresented in international phylogenetic analyses has immediately contributed to global RSV genomic databases with the deposition of all 49 high-quality RSV genomes in the GISAID database. Similar to the SARS-CoV-2 GISAID collaboration paradigm, we strongly recommend Bangladesh’s active participation in regional RSV genomic surveillance networks that enable real-time data exchange between South Asian, Southeast Asian, and Middle Eastern nations.

When interpreting the results of this study, it is important to consider a number of its shortcomings. Low-frequency variations may be undetected or have their prevalence underestimated due to the overall sample size, especially the small number of RSV-B sequences (n=6), which limits the statistical confidence of subgroup-specific estimations. Therefore, any inferences made about the dynamics of RSV-B glycosylation, lineage distribution, or F protein variation should be regarded as suggestive rather than conclusive, and validation in larger, multi-season cohorts is necessary. Sequencing was performed on samples with Ct values ≥25, which favors higher viral loads and may not accurately reflect the range of circulating variety, particularly in mild or early-stage infections. The lack of phenotypic functional assays, such as plaque-reduction neutralization testing and pseudovirus inhibition investigations using locally circulating strains, makes it impossible to assess the therapeutic value of S276N and the reported glycan modifications from sequence data alone. The sampling may overrepresent severe phenotypes and overlook asymptomatic or community-level transmission networks because it only included children under five who reported at sentinel hospital locations with SARI or ILI.

Finally, because the study period does not encompass a full RSV epidemic season, longitudinal surveillance over multiple following seasons will be necessary to distinguish enduring endemic traits from transitory post-pandemic importation patterns in the local genetic landscape.

When genomic monitoring is combined with concurrent clinical, epidemiological, and immunological data, its full potential is realized. In addition to host factors (age in months, gestational age, nutritional status, comorbidities) and immune status (RSV seropositivity, prior vaccination, or monoclonal antibody receipt), we suggest that future RSV WGS studies in Bangladesh prospectively link sequence data to standardized clinical outcome variables, such as oxygen requirement, intensive care unit admission, mechanical ventilation, and case fatality. This linkage enables genotype-phenotype correlation analyses, the assessment of whether specific lineages or mutation profiles are associated with disproportionate disease severity, and the clinical validation layer that transforms genomic observations into valuable clinical and public health information.

## Conclusion

We generated 49 complete RSV genomes (43 RSV-A/ON1, 6 RSV-B/BA9) from Bangladeshi children during the 2024–2025 respiratory season, achieving an 83.1% sequencing success rate via Oxford Nanopore Technology within an established hospital-based surveillance network. The co-circulation of five RSV-A lineages and two RSV-B sub-lineages phylogenetically linked to strains from six continents indicates repeated, independent international introductions rather than a single import event. No resistance mutations were identified at the nirsevimab binding site Ø in either subtype, and all key antigenic sites targeted by licensed vaccines (Arexvy, Abrysvo, mRESVIA) and monoclonal antibodies (nirsevimab, palivizumab) remain intact, supporting the immunological effectiveness of current countermeasures in this setting. RSV-A universally acquired an N-glycosylation site at F-N75, and 67% of RSV-B strains gained a new G protein N-glycan at N256, both representing sites of progressive glycan remodeling that warrant continued surveillance and experimental validation by mass spectrometry. All 49 genomes were deposited in GISAID, contributing to South Asian representation in global RSV phylodynamic databases. These findings provide the evidence base to support nirsevimab and maternal RSV vaccine deployment in Bangladesh and call for the integration of whole-genome sequencing into routine national RSV surveillance.

## Acknowledgments

This research protocol was funded through the generous support of core donors who provide unrestricted funding to icddr,b for its operations and research initiatives. Currently, the Government of the People’s Republic of Bangladesh and Canada are among the key contributors offering this invaluable support, and we sincerely acknowledge and appreciate their commitment to advancing icddr,b’s research efforts.

## Author contributions

Mustafizur Rahman and Mst. Noorjahan Begum conceptualized the study and designed the research framework. Md. Shaheen Alam led all wet laboratory activities, including WGS PCR optimization, library preparation, and data analysis; he also interpreted the results and prepared the initial draft of the manuscript. Yeasir Karim, Rubel Howlader, Riaz Rahman Shanto, and Muhammad Talha carried out laboratory work encompassing diagnostic testing, genotyping, sequencing, and subsequent data analyses. Tanzim Rahman provided valuable assistance with sequence data analysis. Fahmida Chowdhury and Mohammad Jubair offered thorough critical reviews of the article, significantly enhancing its intellectual content. All authors carefully reviewed the manuscript draft and unanimously approved the final version for submission.

